# Phytoene synthase 2 in tomato fruits remains functional and contributes to abscisic acid formation

**DOI:** 10.1101/2021.07.19.452896

**Authors:** Prateek Gupta, Marta Rodriguez-Franco, Reddaiah Bodanapu, Yellamaraju Sreelakshmi, Rameshwar Sharma

## Abstract

In ripening tomato fruits, the leaf-specific carotenoids biosynthesis mediated by phytoene synthase 2 (PSY2) is replaced by a fruit-specific pathway by the expression of two chromoplast-specific genes: phytoene synthase 1 (*PSY1*) and lycopene-β-cyclase (*CYCB*). Though both *PSY1* and *PSY2* genes express in tomato fruits, the functional role of PSY2 is not known. To decipher whether PSY2-mediated carotenogenesis operates in ripening fruits, we blocked the *in vivo* activity of lycopene-β-cyclases in fruits of several carotenoids and ripening mutants by CPTA (2-(4-Chlorophenylthio)triethylamine hydrochloride), an inhibitor of lycopene-β-cyclases. The CPTA-treatment induced accumulation of lycopene in leaves, immature-green and ripening fruits. Even, in *psy1* mutants *V7* and *r* that are deficient in fruit-specific carotenoid biosynthesis, CPTA triggered lycopene accumulation but lowered the abscisic acid level. Differing from fruit-specific carotenogenesis, CPTA-treated *V7* and *r* mutant fruits accumulated lycopene but not phytoene and phytofluene. The lack of phytoene and phytofluene accumulation was reminiscent of PSY2-mediated leaf-like carotenogenesis, where phytoene and phytofluene accumulation is never seen. The lycopene accumulation was associated with the partial transformation of chloroplasts to chromoplasts bearing thread-like crystalline structures. Our study uncovers the operation of a parallel carotenogenesis pathway mediated by PSY2 that provides precursors for abscisic acid biosynthesis in ripening tomato fruits.

**Significance statement:** It is believed that in ripening tomato fruits phytoene synthase 2 that drives carotenoid biosynthesis in leaves is redundant. Contrary to this, we show that in phytoene synthase 1 mutant fruit that is bereft of lycopene, the chemical inhibition of lycopene β-cyclases triggers lycopene accumulation. Our results uncover that phytoene synthase 2 remains functional in ripening fruits and provides precursors for abscisic acid formation in fruits.

## Introduction

In nature, carotenoids provide the color to a wide range of plants, flowers, and fruits and are the most widely distributed pigments (**Schwartz et al., 2008**). Differential accumulation of carotenoids gives diverse hues that attract pollinators and herbivores to the flowers and fruits, thus helping in the pollination and dispersal of seeds. Carotenoid biosynthesis occurs not only in photosynthetic organisms but also in some non-photosynthetic fungi and bacteria (**Walter and Strack, 2011**). In green tissues, they serve as accessory pigments in photosynthesis and protect the reaction center from photooxidation (**Müller et al., 2001; Robert et al., 2004**). Carotenoids also serve as a precursor for several volatiles, responsible for the fruits’ taste and aroma **(Baldermann et al., 2010**).

Carotenoid biosynthesis most prominently occurs in chloroplasts, where carotenoids play a key role in photosynthetic light absorption (**Niyogi, 1999**). A multi-pronged approach consisting of analysis of mutants, chemical inhibitors, and labeling studies identified the biochemical steps leading to carotenoid synthesis and degradation (**Spurgeon, 1983**). Above studies established that in higher plants, geranylgeranyl pyrophosphate, the first committed precursor for carotenoid synthesis, derives from the plastid-localized methylerythritol 4-phosphate pathway. While chlorophylls accumulation/synthesis obligatorily needs carotenoids, carotenoids biosynthesis is not coupled to chlorophylls. Carotenoids are also present in non-photosynthetic organs of plants such as flowers and fruits. Among fruits, wide color variations, and shorter time duration needed to transit to a fully colored fruit, makes tomato an ideal system for deciphering molecular-genetic mechanisms regulating carotenoid biosynthesis.

The first rate-limiting step of carotenoid biosynthesis involving condensation of two geranylgeranyl pyrophosphate molecules is catalyzed by phytoene synthase (PSY), generating 15-cis phytoene, a colorless compound (**Fraser et al., 2002**). While Arabidopsis has a single *PSY* gene, in tomato, PSY was triplicated with neofunctionalization of additional genes during evolution. Among these genes, the fruit-specific carotenoid biosynthesis is mediated by a chromoplast-specific paralog of *PSY*, namely *PSY1*. The other two *PSY* genes also show tissue-specific expression, with *PSY2* mediating leaf-specific carotenoids biosynthesis, whereas expression of *PSY3* is limited to roots under stress conditions **(Fraser et al., 1999**; **Kachanovsky et al., 2012; Fantini et al., 2013**).

In conformity with the role of *PSY1* in regulating fruit-specific carotenoids biosynthesis, a loss-of-function mutation-*yellow flesh* (locus *r*), causes a severe reduction in carotenoids in ripe tomato fruits. The carotenoids reduction in the *r* mutant can be recovered by transgenic-overexpression of a wild-type copy of *PSY1* (**Fray and Grierson, 1993**). The colorless phytoene is converted by phytoene desaturase (PDS), through a series of desaturations, to 15-*cis* phytofluene and 9,15,9’-tri-*cis*-ζ-carotene, which undergoes isomerization to form 9,9’-di-*cis*-ζ-carotene by ζ-carotene isomerase (ZISO). Alike *PSY1, zeta* (*z*^*2803*^) mutation in *ZISO* terminates carotenoid biosynthesis at ζ-carotene in tomato fruit **(Kachanovsky et al., 2012**).

Di-*cis*-ζ-carotene is desaturated by ζ-carotene desaturase (ZDS) to form 7,9,9’-tri-*cis*-neurosporene and 7,7’,9,9’-tetra-*cis*-lycopene. Isomerization of tetra-*cis*-lycopene to all-*trans*-lycopene is catalyzed by a carotenoid isomerase (CrtISO) encoded by the *tangerine* (*t*) locus of tomato. Two *tangerine* mutant alleles have been reported: *tangerine*^mic^, affected by a deletion of the *CRTISO*, and *tangerine*^3183^, with impaired *CRTISO* expression (**Isaacson et al., 2002**). The carotenoid isomerization mediated by ZISO and CRITSO can also be directly carried out by the light (**Isaacson et al., 2002**; **Fantini et al., 2013**).

Using VIGS, **Fantini et al. (2013**) delineated the relative contribution of *PSY1, PDS, ZDS, ZISO, CrtISO, CrtISO*-Like1, and *CrtISO*-Like2 to tomato fruit carotenogenesis. *PSY1* and *CrtISO*-silenced fruits displayed a phenotype similar to *yellow flesh* and *tangerine* mutants, respectively. Consistent with light acting as an effector for carotenoid isomerization, the light exposure restored lycopene biosynthesis in *ZISO*-silenced fruits. All-*trans*-ζ-carotene was detected in *ZDS*-silenced fruits but not in *CrtISO*-Like1-/*CrtISO*-Like2-silenced fruits. These studies suggested isomerization to all-*trans*-lycopene as an additional regulatory step in carotenoid biosynthesis.

The mode of cyclization of all-*trans-*lycopene acts as a decisive point in controlling the flux of carotenoids to the α-carotene or the β-carotene branch. The conversion of lycopene to α-carotene or β-carotene is governed by lycopene ε- and β-cyclases (LCYE and LCYB), respectively. LCYE adds an ε-ring to all-*trans* lycopene to form a monocyclic δ-carotene, which gets β-cyclized on the opposite end by LCYB to form α-carotene. The conversion to β-carotene is mediated by the addition of one β-ring to all-*trans*-lycopene to form γ-carotene, which is converted to β-carotene by the addition of one more β-ring by LCYB (**Cunningham et al., 1996**). Similar to *PSY1*, during evolution, *LCY* first diverged to *LCYE* and *LCYB*, and then to chloroplastic *LCYB1, LCYB2*, and chromoplastic *CYC-B* (**Mohan et al., 2016**).

Studies in tomato revealed that carotenoid pathway genes display developmental and organ-specific regulation during fruit development (**Lois et al., 2000**). Though *PSY2* seemingly does not contribute to carotenogenesis, it expresses during tomato ripening, albeit at much lower levels than *PSY1* (**Giorio et al., 2008; Kilambi et al., 2021**). Similarly, *LCY-E, LCYB1*, and *LCYB2* expression continue along with *CYC-B* in ripening fruits (**Kilambi et al., 2021**). It remains to be determined whether these genes are redundant, or have a function hitherto not reported, due to the paucity of mutations in these genes. The absence of mutations for critical genes is generally overcome by the chemical inhibition of the genes. Consistent with this, inhibition of PDS by Norflurazon blocks carotenogenesis and bleaches photosynthetic tissue due to photooxidation of chloroplasts (**Oelmüller and Mohr, 1986; Chamovitz et al., 1991**). The inhibitors targeting PDS have been used as a tool to study the up/downstream carotenogenic gene expression in different tissues (**Simkin et al., 2000**).

Similar to PDS, the activity of lycopene cyclases is sensitive to 2-(4-Chlorophenylthio) triethylamine hydrochloride (CPTA) (**Coggins et al., 1970**). The CPTA application blocks the conversion of lycopene to α-carotene or β-carotene, preventing the flux into these pathways. Consequently, the carotenoid biosynthesis in CPTA-treated tissue is terminated at lycopene, leading to ectopic lycopene accumulation even in photosynthetic tissues (**Knypl, 1969**).

Though in tomato fruits both *PSY1* and *PSY2* genes express, the functional role of PSY2 is not known. To uncover its role, we examined carotenogenesis in fruits of several *psy1*, and carotenogenesis mutants of tomato treated with or without CPTA. To complement this, we also treated tomato ripening mutants *green flesh, rin*, and *nor* with CPTA. We report that contrary to the assumption that PSY1 is the sole active enzyme in tomato fruits, PSY2 is also functional and contributes to abscisic acid formation in fruits. Seemingly PSY2-mediated pathway does not contribute to the formation of the fruit-like complement of carotenoids and is seen only because CPTA-treatment terminated it at lycopene.

## Results

CPTA is known to cause lycopene accumulation in a wide range of plant tissues (**Coggins et al., 1970**), including in yellow lutescent tomatoes, which are devoid of lycopene (**Jen and Thomas, 1977**). It is believed that during tomato fruit ripening, leaf-specific carotenogenesis is replaced by fruit-specific carotenogenesis, thus fruits have carotenoids complement different from photosynthetic leaves. Using CPTA as a tool, we investigated whether the PSY2 which is a key enzyme for leaf-specific carotenogenesis continues to operate in tomato fruit and its role thereof in fruits.

### Effect of CPTA on carotenoid producing *E. coli* strains

To check whether CPTA indeed blocks the activity of lycopene cyclases, we first studied the effect of CPTA (1 mM) on *E. coli* expressing different carotenoid pathway genes viz., ζ-carotene (pAC-ZETA), lycopene (pAC-LYC), ε-carotene (pAC-EPSILON), β-carotene (pAC-Beta-At) and pAC-85b (produces β-carotene when complemented with functional phytoene synthase). The addition of CPTA to pAC-EPSILON, pAC-ZETA, and pAC-Beta-At cultures produced lycopene as the main carotenoid evidently by inhibiting *LCYE* and *LCYB*, respectively. In contrast, CPTA addition to the pAC-85b, pAC-ZETA, and pAC-LYC did not affect the carotenoid content (**Figure S1**). These experiments reiterated that CPTA specifically acts on lycopene cyclases.

### Effect of CPTA on leaf tissue

We then examined the efficacy of CPTA on lycopene accumulation by irrigating tomato seedlings with different amounts of CPTA. The retardation of plant growth, and bleaching of leaves at margins, due to reduction of photoprotective carotenoids level indicated efficient translocation of CPTA to leaves (**Figure 1A**). CPTA has a high solubility in the lipids, thus can cross cell membranes (**Poling et al., 1975**) and thereby can enter the xylem stream. The lycopene accumulation was observed in >100 μM CPTA-treated leaves. The bleached portion harvested from 500 μM CPTA-treated leaves showed a massive accumulation of lycopene (35 μg/gm FW) (**Figure 1B**). Conversely, the level of β-carotene declined with increasing concentrations of CPTA. Interestingly, CPTA induced the accumulation of γ-carotene and δ-carotene, which were absent in control leaves. Notably, the α-carotene level rose in 100 μM CPTA-treated leaves and then declined. Notably, though CPTA triggered lycopene accumulation, no other carotenoid upstream of lycopene such as phytoene or phytofluene accumulated.

**Figure 1.**
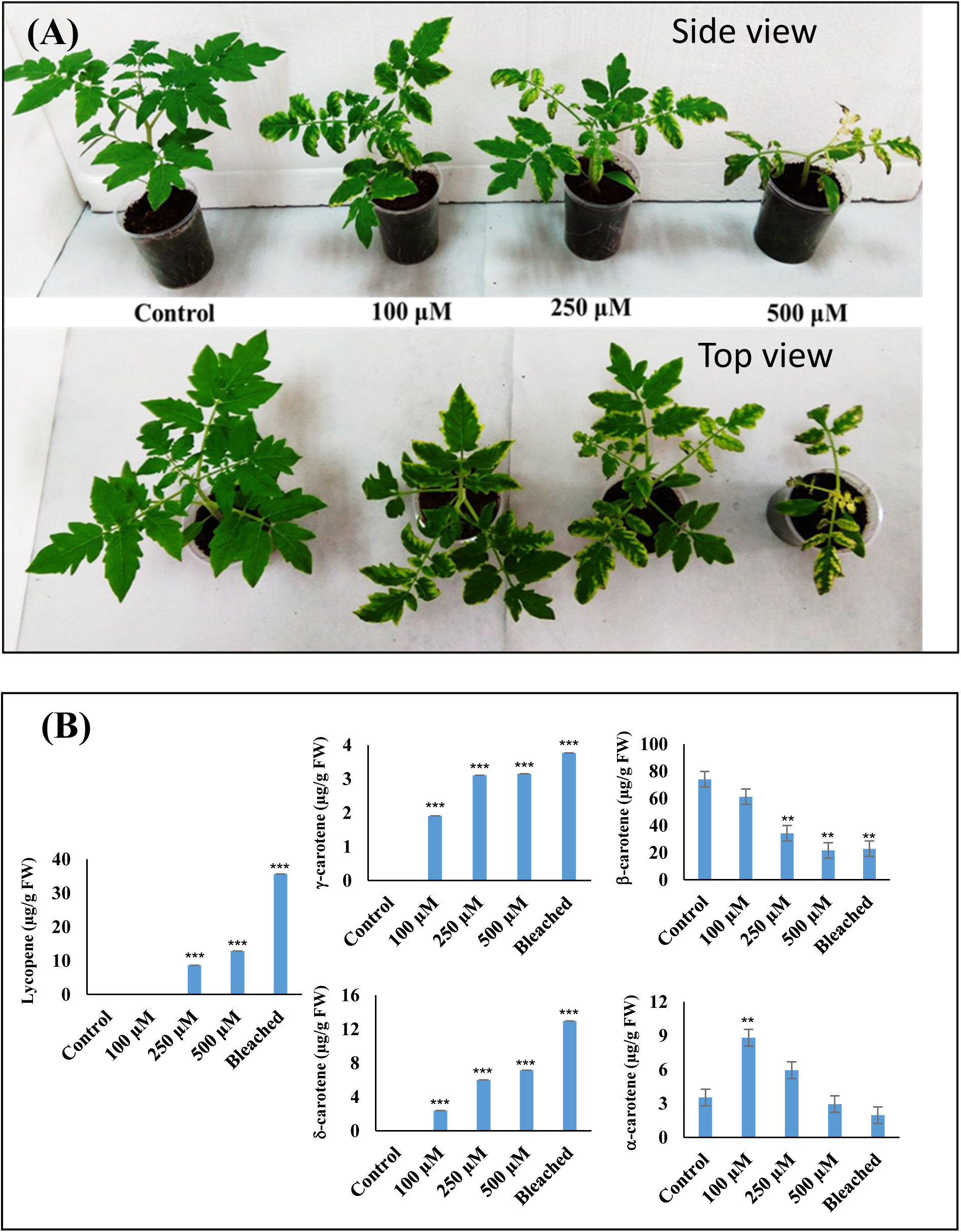
Effect of different concentrations of CPTA on the growth of Arka Vikas plants. (**A**) The side and top view of the plant phenotype (**B**) Carotenoid profiling from leaves of the same plants. The bleached portion was collected only from 500 μM CPTA-treated leaves. Seedlings for two weeks were raised using water irrigation. After that, seedlings were irrigated with 50 ml of desired CPTA concentration, and the same repeated after a one-week interval. Leaves were collected after two-week of CPTA treatment. The CPTA effect was more intense towards plants’ apex, as it blocks the formation/accumulation of carotenoids in younger leaves. Note bleaching of margins and reduction of growth due to loss of photosynthetic competence. (n=3, * p ≤ 0.05, ** p ≤ 0.01, *** p ≤ 0.001, p-values are calculated with respect to control).

### Chronological study of CPTA effect

We next examined the effect of CPTA on fruits. To find an optimal amount for experimentation, we injected CPTA solutions of varying molarity (1, 10, 50, 100, 200 mM) into fruits. The fruits ranged from different immature-green (IG) to mature-green (MG) stages of development. Post-injection fruits were visually monitored for 15 days. The IG-fruits developed pinkish color, whereas MG-fruits developed deeper-red color than controls. However, 200 mM CPTA triggered fruit deterioration, partially in MG but wholly in IG (**Figure S2**). Considering this, we selected 100 mM CPTA for further experiments.

To investigate the CPTA effect in detail, we injected CPTA into four fruits of the same age, while controls were injected with water. Treated fruits were individually profiled for carotenoids. We used three fruit developmental stages, which consisted of two IG stages based on the fruit size, IG1 being the smallest, IG2, and the MG stage. Post-injection visual monitoring of detached control fruits revealed that IG1 remained green, IG2 turned orangish, and MG attained red color. However, in CPTA-treated fruits, IG1 attained pink, IG2 red, and MG developed deep red coloration (**Figure S3, S4**). The carotenoid profile correlated with the fruit color; CPTA-treated fruits had a higher accumulation of all-trans lycopene than the control. The orangish color of IG2 control fruits resulted from the accumulation of β-carotene during incubation. Conversely, in CPTA-injected fruits, β-carotene and lutein declined in a pattern opposite to lycopene (**Figure S3**). Notably, while the control IG1 and IG2 fruits did not accumulate phytoene and phytofluene, CPTA-induced phytoene and phytofluene accumulation in parallel to lycopene.

CPTA-treatment also altered the ultrastructure of plastids. Control MG and green-colored IG1 and IG2 fruits had a chloroplast-like structure with stacks of the thylakoid membrane. In control orange-colored IG2 fruit, most grana stacks were dissolved, and the plastoglobule number increased. In CPTA-treated fruits, crystalline thread-like structures, characteristically associated with lycopene accumulation, appeared along with fewer plastoglobules. Though lycopene threads were also visible in water-treated ripened MG fruit, these were fewer than in CPTA-treated fruits (**Figure S4**).

To decipher the time course of CPTA-induced carotenoid accumulation, fruits injected at the MG stage were harvested daily and transversely cut to visually monitor the color. The red coloration appeared on the fourth day and became intense from the fifth day onwards. Remarkably, the columella turned red in treated fruits compared to untreated tomato fruits (**Figure S5**). Carotenoid profiling of the same CPTA-treated fruits revealed a sustained increase in phytoene and phytofluene from day four onwards, a higher lycopene level from day five onwards, and a block in the accumulation of β-carotene (**Figure S6**).

### CPTA induces lycopene accumulation in mutants impaired in carotenoid accumulation

We next examined whether CPTA can induce lycopene in tomato mutants/cultivars compromised in carotenoid accumulation or fruit ripening. These mutants accumulate no or little lycopene, and ripened fruits remain green or orange in color. CPTA was injected in MG fruits of *r* and V7 (*psy1*-mutants) (**Mickey, 2013**), *yellow oxheart* (*crtiso-*mutant) (https://www.heritagefoodcrops.org.nz/table-of-tomatoes-containing-tetra-cis-lycopene/), IIHR 2866 (high β-carotene line) (**Kavitha et al., 2014**), *rin* (*MADS*-box transcription factor-mutant), *nor* (*NAC*-domain transcription factor-mutant) (**Barry, 2014**), and green-flesh (*gf, SGR* gene-mutant) (**Barry et al., 2008**). Alike Arka Vikas (control variety), CPTA-injected mutant fruits accumulated lycopene, while the amount of β-carotene and xanthophylls declined (**Figure 2**). CPTA-treated fruits of V7, *r, nor*, and *rin* mutants did not show accumulation of phytoene and phytofluene; however, the levels of these two compounds increased in the treated *gf* mutant and the IIHR 2866 line. The *yellow oxheart* mutant showed no perceptible CPTA induction of lycopene, probably due to the loss of the CRITSO function. The absence of lycopene accumulation in *yellow oxheart* mutant could also be due to the lack of light penetration beyond the outer pericarp, which is needed for cis/trans isomerization of prolycopene to lycopene in fruits (**Issacson et al., 2002**).

**Figure 2:**
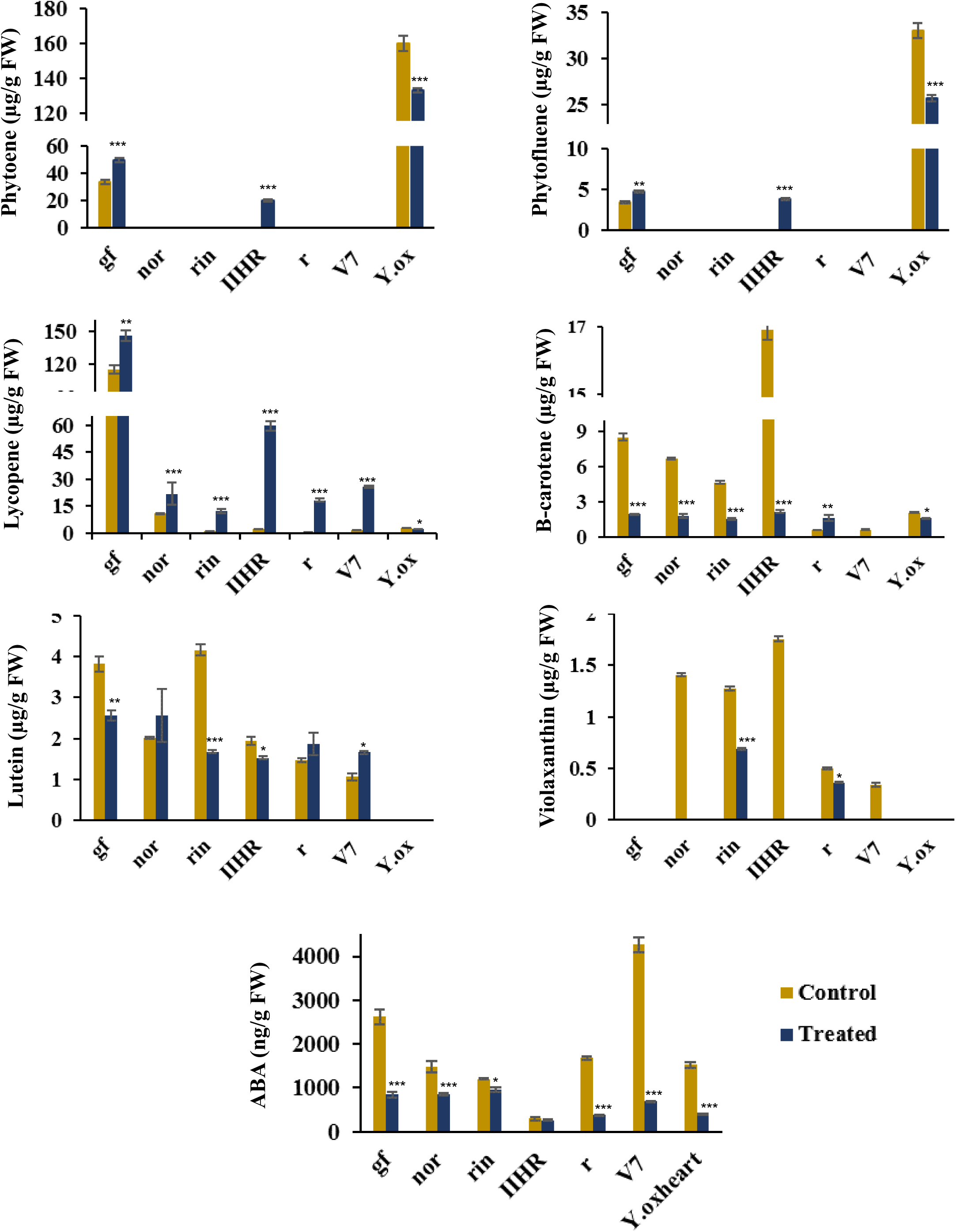
Carotenoid and ABA profiling in fruits of different tomato lines injected with CPTA. The fruits were injected with CPTA/water at the mature green stage, and the carotenoids and ABA levels were measured after 12 days from injections. The following tomato lines were used: *greenflesh*-*gf, nor, rin*, IIHR2866, *r, V7*, and *Yellow oxheart* (*Y*.*ox*). (n>=4, * p ≤ 0.05, ** p ≤ 0.01, *** p ≤ 0.001, p-values are calculated for the treated samples with respect to respective control).

Interestingly, CPTA-injected cut-halves of *V7, r, rin, nor, gf*, and IIHR 2866 fruits showed varying shades of muddy-red color distributed across the fruit, including columella (**Figure 3**). Consistent with loss of CRTISO function and absence of light penetration in deeper fruit tissues (**Issacson et al., 2002**), CPTA-treated *yellow oxheart* fruits exhibited no color change in the placenta. Nonetheless, the *yellow oxheart* fruits showed a tinge of red color in the outer pericarp. It can be construed that in *yellow oxheart* mutant CPTA-induced lycopene formation was limited to the region of light penetration. We then examined whether CPTA-induced decrease in xanthophylls also causes a reduction in abscisic acid (ABA), a downstream product derived from xanthophylls. Hormonal profiling of CPTA-treated fruits showed a massive decline in ABA levels compared to respective controls (**Figure 2**). The ABA decline was very prominent in *psy1* (*V7* and *r*) and *crtiso* (*yellow oxheart*) mutants.

**Figure 3:**
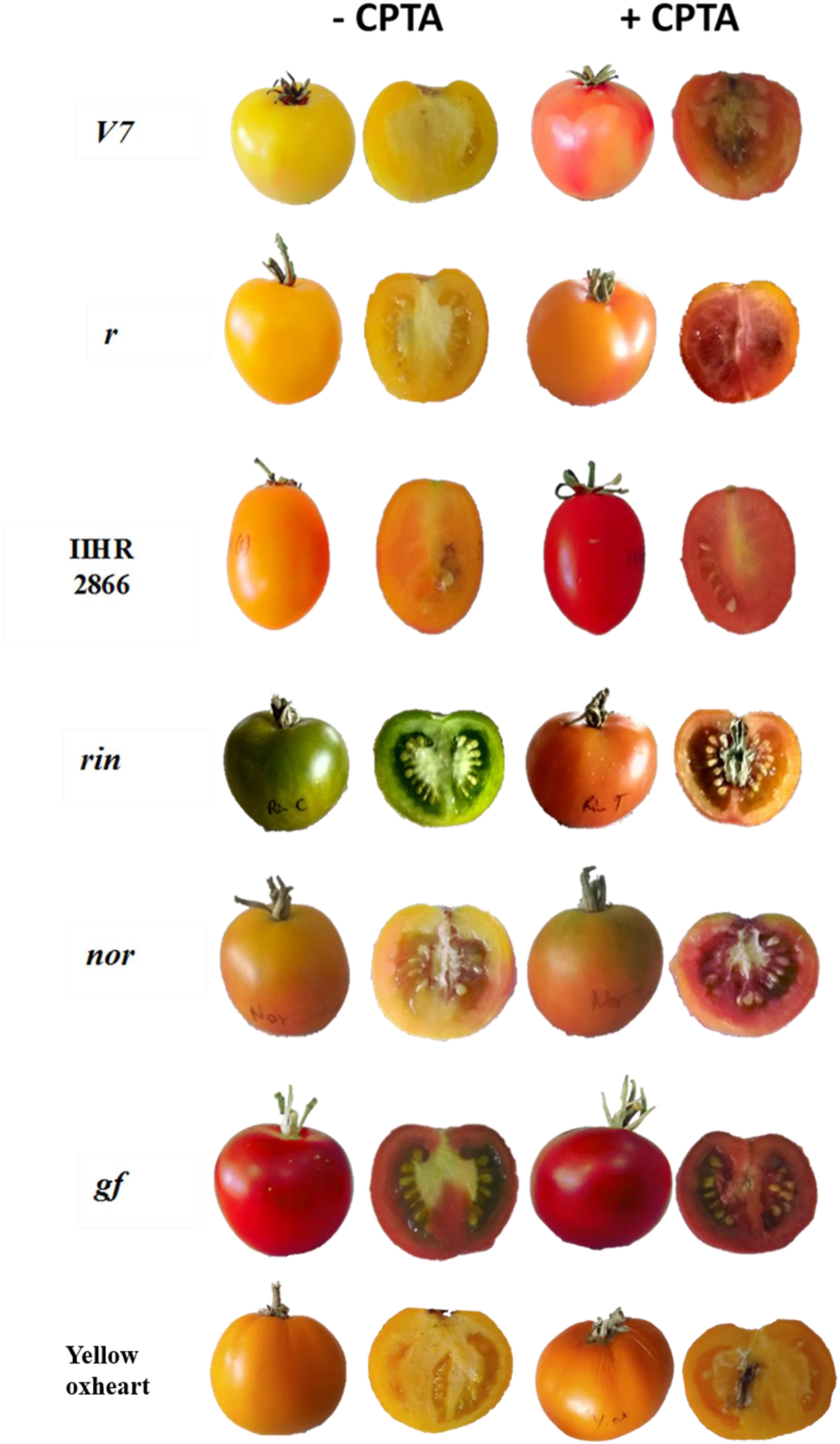
The appearance of red coloration in tomato fruits injected with or without CPTA post-12 days of injection. The following tomato lines were used: *greenflesh*-*gf, nor, rin*, IIHR2866, *r, V7*, and *Yellow oxheart* (*Y*.*ox*). Note in IIHR 2866, the lycopene accumulation results due to inhibition of β-carotene formation in CPTA-treated fruits.

### CPTA does not influence gene expression levels of carotenoid pathway genes

To ascertain whether CPTA treatment also altered carotenoid pathway genes’ expression, we examined transcript levels of genes belonging to methylerythritol 4-phosphate and carotenoid biosynthesis pathways (**Figure S7**). The genes belonging to the methylerythritol 4-phosphate pathway, 1-deoxy-D-xylulose 5-phosphate synthase (*DXS*), 1-Deoxy-D-xylulose 5-phosphate reductoisomerase (*DXR)*, and geranylgeranyl pyrophosphate synthase (*GGPPS2)* showed variation in expression level across the mutants. However, in the majority of the mutants, these genes were significantly not affected by the CPTA treatment. Similarly, genes belonging to chromoplast specific (fruit)-viz., *PSY1, CYCB*, and chloroplast-specific (leaf) carotenoid pathways *PSY2, LCYB1, LCYB2, LCYE* were not altered in the treated fruits. The same was observed for common genes of the carotenoid biosynthesis pathway *ZDS, ZISO, CRTISO*, and carotenoid cleavage dioxygenase (*CCD)* genes, *CCD1A, CCD1B, CCD4A, CCD4B, CCD7*, and *CCD8*.

### Volatile analysis of mutants impaired in carotenoid accumulation

Many volatiles of tomato are derived from the breakdown of carotenoids. Thus, a shift in carotenoid composition also alters the volatile profiles. Considering the CPTA-altered carotenoid composition, we monitored the volatile profiles of control and treated fruits. The major volatiles released from the breakdown of lycopene and other noncyclic tetraterpenoids are geranial, neral, 6-methyl-5-hepten-2-one, and (E,E)-pseudoionone. In contrast, β-Ionone is prominent in fruits containing β-carotene (**Lewinsohn et al., 2005**). Consistent with lycopene accumulation, CPTA-treated fruits had higher amounts of volatiles derived from noncyclic carotenoids. Conversely, β-Ionone derived from the β-carotene massively declined in CPTA-treated fruits of IIHR2866 (**Figure 4**). Broadly, the volatile profiles reflected the carotenoid composition of the respective treated and untreated fruits. The volatile composition also signified the termination of carotenogenesis in CPTA-treated fruits at lycopene.

**Figure 4:**
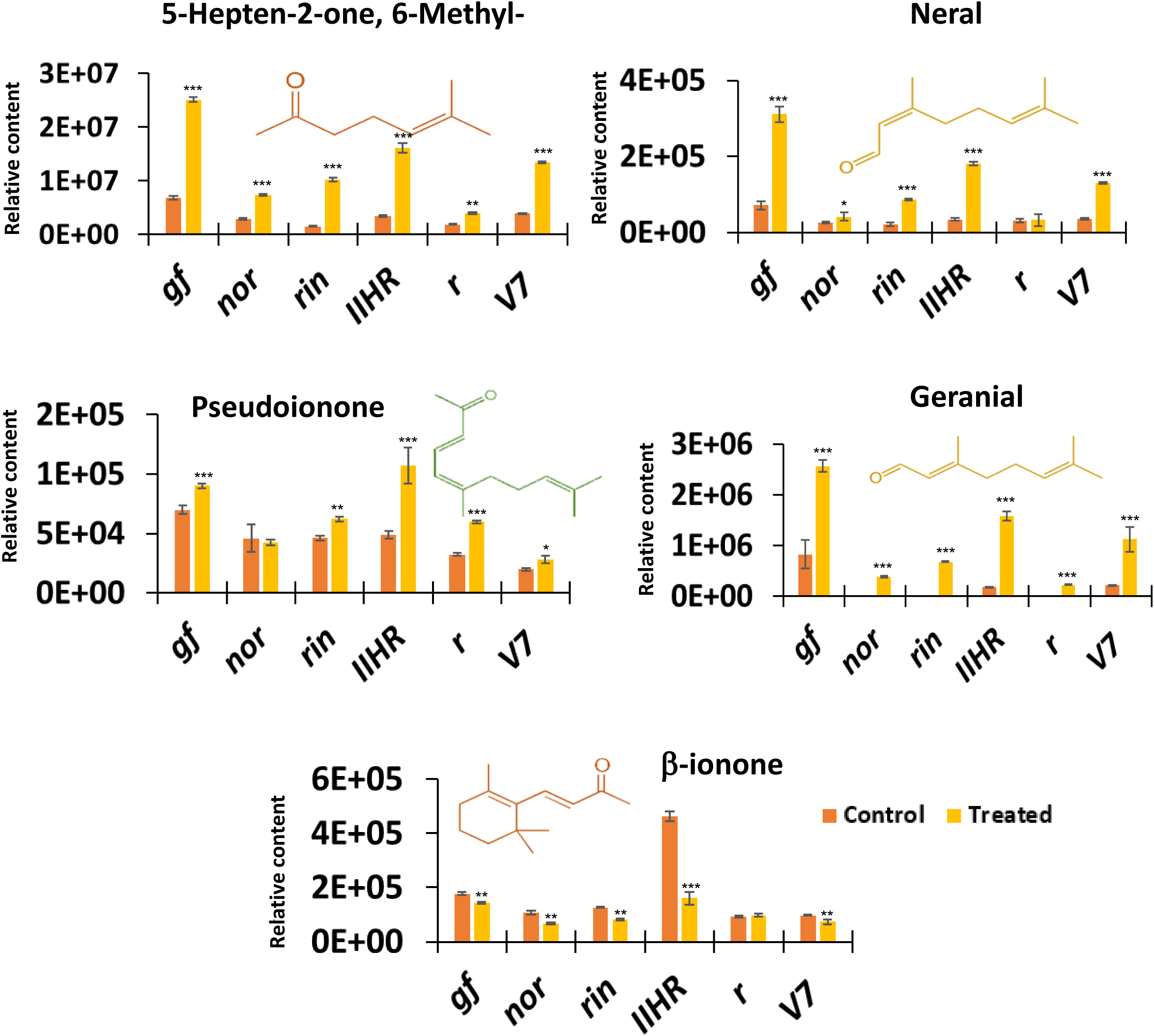
Carotenoid-derived volatiles in CPTA-injected *V7, r*, IIHR 2866, *rin, nor*, and *gf* fruits. Fruits were injected with CPTA at the mature green stage, and profiling was done after 12 days of treatment. (n>=4, * p ≤ 0.05, ** p ≤ 0.01, *** p ≤ 0.001, p-values are calculated for the treated samples with respect to respective control).

### CPTA-triggered accumulation of lycopene-crystalloids in plastids

To check whether CPTA-triggered chromoplast formation, the pericarps of control and CPTA-treated fruits were examined for autofluorescence. On light excitation, chlorophyll-rich plastids emit red, carotenoids-rich plastids emit green, and intermediate plastids emit orangish/yellow fluorescence due to the merging of green and red fluorescence. The *rin, r*, and *nor* plastids emitted both green and red fluorescence at 500-550 nm and 650-700 nm, respectively. The merged images displayed yellowish-orange color, confirming the presence of both chlorophylls and carotenoids. In contrast, *V7* plastids emitted only green fluorescence indicating the exclusive accumulation of carotenoids (**Figure S8**). Consistent with the emission of carotenoids-specific autofluorescence, the ultrastructure of *V7, r, rin, nor, gf*, and IIHR 2866 plastids showed varying degrees of chloroplast to chromoplast conversion. of lycopene crystalloids (**Figure 5**). Taken together, CPTA-triggered the ectopic accumulation of lycopene and chromoplast transformation in the above mutants.

**Figure 5:**
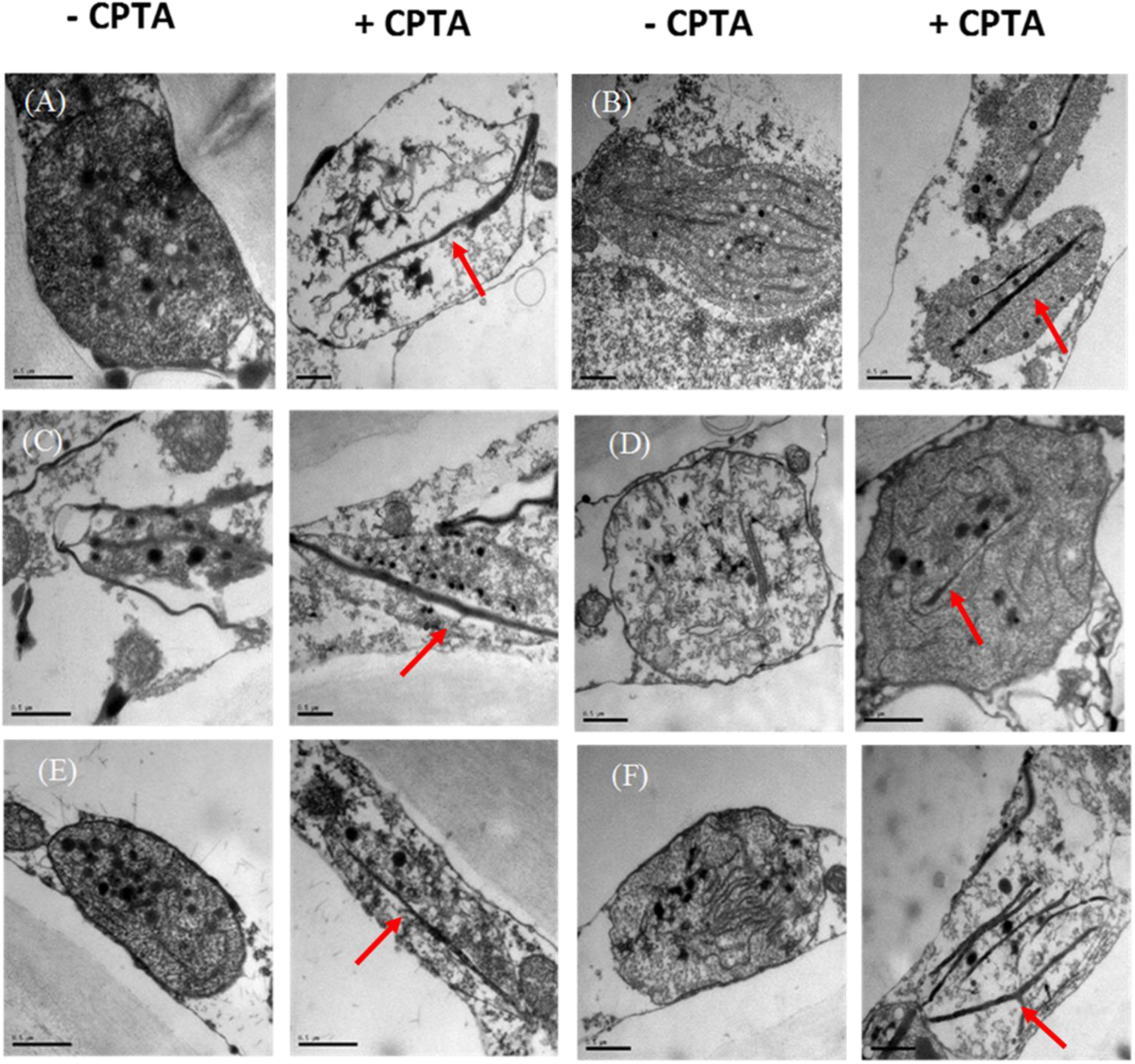
Ultrastructure of plastids of fruits treated with or without CPTA **(A)** *V7*, (**B**) *rin*, (**C**) *gf*, (**D**) *r*, (**E**) *nor* and (**F**) IIHR 2866. Note formation of distinct lycopene threads (marked by red arrow) in plastids of CPTA-treated fruits, signifying partial conversion of plastids into chromoplast. The bar in the figure is equal to 0.5 µm.

### CPTA induces lycopene accumulation in *yellow flesh* mutant fruits in dark

The conversion of phytoene to lycopene requires a set of enzymes that carry out isomerization and desaturation. The isomerization step carried out by *ZISO* and *CRTISO* can also be performed by light. Between *ziso* and *crtiso* mutants, light can efficiently restore carotenoid biosynthesis in *ZISO*-silenced fruits, compared to *CRTISO*-silenced fruits (**Fantini et al., 2013**). Taking advantage of this, we treated fruits of *r, r*^*3756*^ (mutant line with yellow-flesh phenotype having an early stop codon in the *PSY1* gene) (**Kachanovsky et al., 2012**), and *ziso* mutant at MG stage with CPTA. The fruits were kept either in total darkness or in dark/light cycle conditions. In light, CPTA-treated fruits of *r*, and *r*^*3756*^ mutants accumulated lycopene, but not phytoene and phytofluene. Whereas, *ziso* mutant accumulated lycopene along with phytoene and phytofluene.

Similar to light, the CPTA-treated dark-incubated *r*, and *r*^*3756*^ fruits accumulated lycopene, albeit, at the reduced magnitude, but not phytoene and phytofluene. Predictably, the dark-incubated *ziso* mutant completely lacked lycopene (**Figure 6**) but accumulated phytoene and phytofluene. The above results are in conformity with the view that while PSY2-mediated carotenogenesis operates in *r*, and *r*^*3756*^ tomato fruits, nonetheless above carotenogenesis pathway needs functional *ZISO* at least in the darkness.

**Figure 6:**
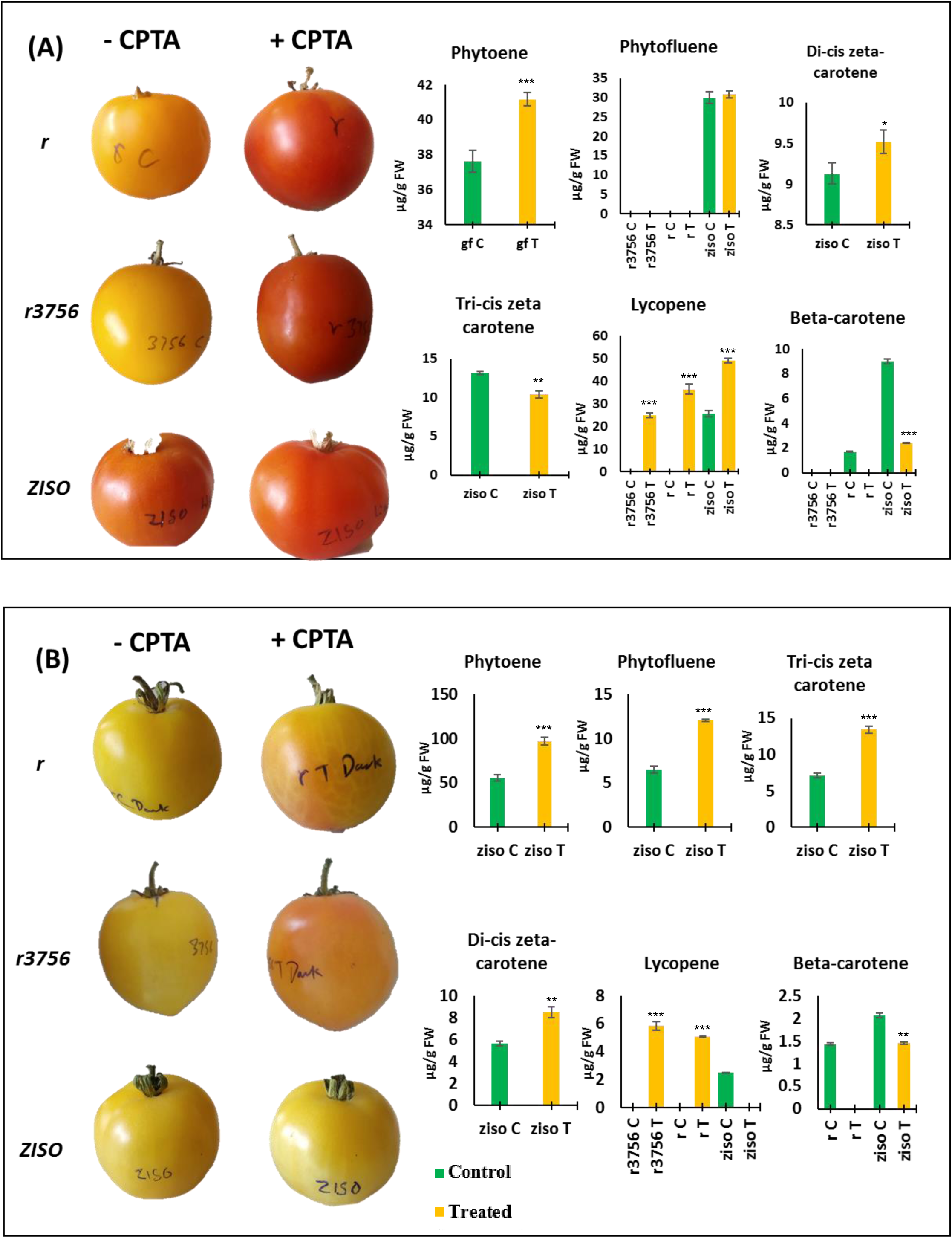
Influence of light and dark treatments on the accumulation of lycopene in *r, r*^*3756*^ and *ziso* mutant fruits. The detached fruits after water-(**C**) or CPTA-injection (**T**) were incubated in light (16h-light/8h-dark cycle) (**A**) or in total darkness (**B**). Note the absence of lycopene formation in CPTA treated dark-grown *ziso* fruits (n>=3, * p ≤ 0.05, ** p ≤ 0.01, *** p ≤ 0.001, p-values are calculated for the treated samples with respect to respective control).

## Discussion

The biosynthesis of carotenoids, including its precursor geranylgeranyl pyrophosphate derived from the methylerythritol 4-phosphate pathway, is exclusively localized in plastids. In consonance with its role in the photoprotection of photosynthetic centers, carotenoid biosynthesis in chloroplasts is coupled with chlorophylls to maintain the chlorophyll/carotenoids stoichiometry **(Sun et al., 2018)**. In tomato fruits, the transition to ripening uncouples carotenoids biosynthesis from chlorophylls leading to the transformation of chloroplasts into carotenoid-rich chromoplasts. The above transition in tomato fruits also involves a shift in the first committed step for carotenoid biosynthesis mediated by phytoene synthase (PSY). The onset of ripening upregulates the expression of fruit-ripening-specific *PSY1*, while leaf-specific *PSY2* continues to express at subdued levels (**Fraser et al., 1994; Fraser et al., 1999; Giorio et al., 2008; Kilambi et al. 2013**). We show that PSY2 is not redundant in chromoplasts; it continues to function in ripening fruits and provides precursors for ABA synthesis in fruits.

### CPTA-treated leaves accumulate lycopene but not phytoene and phytofluene

To ascertain that CPTA-treatment did not influence the PSY2 activity and intermediary enzymes leading to lycopene formation, tomato seedlings were irrigated with CPTA. Consistent with CPTA being a specific inhibitor of lycopene cyclases, CPTA-treated leaves accumulated lycopene. The marginal bleaching of leaves is consistent with the release of CTPA from the xylem stream at leaf margins (**Shapira et al., 2009**). The growing leaves expand at margins, where newly formed cells synthesize chlorophylls, and CPTA-blockage of xanthophylls accumulation causes bleaching. The accumulation of lycopene in leaves indicated that PSY2 mediated carotenoid synthesis in leaves was terminated by CPTA at lycopene due to inhibition of lycopene cyclases.

Notably, CPTA-treated leaves accumulated none of the intermediate carotenes upstream to the lycopene, such as phytoene and phytofluene, while showed an increase in levels of γ-carotene and δ-carotene. The accumulation of γ-carotene and δ-carotene, along with lycopene, is consistent with the notion that CPTA is less effective in blocking the first lycopene cyclization reaction than the second reaction (**Young et al., 1989; La Rocca et al., 2007**). In leaves, CPTA more effectively blocked the β-carotene branch than the α-carotene branch, as evidenced by the increase in lutein and α-carotene in treated leaves. The reduced growth of plants may come from a multifold effect of CPTA, as in addition to photobleaching, CPTA may have influenced the levels of growth regulatory apocarotenoids **(Hou et al., 2016**).

### Lycopene accumulation induces precocious chromoplasts-like plastids in immature green fruits

An ideal way to decipher the role of PSY2 in tomato fruits would be to use a *PSY2* mutant. Lamentably, there are no reported tomato *PSY2* mutants. The sole reported phytoene synthase mutant in Chlamydomonas, namely *light-sensitive 1*, lacks plastid ultrastructure and can be only heterotrophically maintained (**Inwood et al., 2008**). The mutations in PSY2 may be lethal as it will lead to total loss of photosynthesis, thus so far not been recovered in mutagenesis screens. Though phytoene desaturase, the next enzyme in the pathway, can be specifically blocked by Norflurazon (**Römer et al., 2000**), its potential is limited, as phytoene accumulation may inhibit PSY activity (**Simkin et al., 2003; Campisi et al. 2006**). The CPTA that blocks lycopene cyclases proffers a better alternative, as it allows carotenoid biosynthesis to proceed up to lycopene, allowing the normal functioning of upstream enzymes.

The precocious formation of lycopene in CTPA-treated immature green fruits is in conformity with the blockage of lycopene cyclases, irrespective of the fruit’s developmental stage. The precocious formation of chromoplasts-like plastids in CPTA-treated immature-green fruits also indicated a linkage between lycopene accumulation and chromoplast formation. Alternatively, the appearance of chromoplasts-like plastids may ensue from photooxidative loss of thylakoids accompanied by accumulation of lycopene in plastid stroma (**La Rocca et al., 2007**). The chromoplast like structure formation in IG1 and IG2 fruits can also be caused by the ectopic formation of phytoene triggered by CPTA (**Llorente et al., 2020; Andersen et al., 2021)**

Post-MG stage, the carotenogenesis pathway in tomato fruits is modified to stimulate lycopene and β-carotene accumulation. The reduction in β-carotene levels by the CPTA treatment is consistent with the termination of carotenogenesis at the lycopene step. Likewise, the increase in phytoene and phytofluene in treated fruits seems to ensue from the blockage of carotenogenesis. The increase in phytoene and phytofluene in MG fruits is also consistent with the notion that CPTA does not inhibit enzymes upstream to the lycopene, at least PSY and PDS activity (**Al-Babili et al., 1999**). Considering that, unlike leaves and control fruits, CPTA-treated IG1 and IG2 fruits accumulate phytoene and phytofluene, it can be surmised that the above accumulation ensues from the action of PSY1. This view is consistent with the observation that *PSY1* accumulation is triggered in the immature green fruits, much before the onset of ripening (Lois et al., 2000).

### PSY1 and PDS activity seems to be not affected by CPTA

Considering that *yellow oxheart*, mutated in *CRITSO*, hyperaccumulates phytoene and phytofluene, and CPTA does not affect these intermediates supports the view that CPTA does not block PSY and PDS enzymes. Consistent with this, the *green flesh* mutant with no mutations in any carotenoid biosynthesis genes accumulates high phytoene and phytofluene levels, along with lycopene and β-carotene. Conversely, a high β-carotene tomato cultivar IIHR2866 accumulates only traces of lycopene, most likely due to the efficient conversion of lycopene to β-carotene. CPTA-stimulated lycopene accumulation and reduction in β-carotene in *green flesh* and IIHR2866 are in conformity with the view that the CPTA effect is restricted to lycopene cyclases. This view is also consistent with the effect of CPTA on bacteria expressing carotenoids biosynthesis genes, wherein lycopene accumulation and β-carotene loss are observed due to inhibition of lycopene cyclases.

### CPTA treatment unmasks carotenoids biosynthesis in fruits that is independent of PSY1

The onset of ripening in tomato leads to overexpression of fruit-specific PSY1, relegating leaf-specific PSY2 to presumably ineffectual function. Considering that *PSY1* mutants such as *V7, r* (**Fray and Grierson 1993**), and *r*^*3756*^ (**Kachanovsky et al., 2012**) fail to accumulate lycopene in ripe fruits, the lycopene biosynthesis is blocked in these fruits due to nonfunctional PSY1. An alternative blockage is seen in non-ripening mutants of tomato viz. *rin* and *nor*, encoding transcription factors, where lycopene does not accumulate due to the absence of *PSY1* expression (**Ito et al., 2017; Thompson et al. 1999**). In all these mutants, the PSY1-mediated carotenogenesis leading to lycopene seems to be blocked, as these mutants do not accumulate phytoene or phytofluene.

Interestingly, though *V7, r, rin*, and *nor* do not accumulate lycopene, they have β-carotene. It can be construed that β-carotene is a remnant of photosynthetic carotenoids from mature green fruits that are retained in these mutants. This view is consistent with the analysis of leaf and mature green fruits carotenoids where phytoene and phytofluene do not accumulate but have β-carotene and other xanthophylls. Notwithstanding the above considerations, it is equally plausible that post-MG phase, β-carotene present in fruits is also synthesized by a pathway that does not involve PSY1. In that situation, it could be that PSY2 may still be operational in chromoplasts, albeit at a subdued level. The accumulation of lycopene in CPTA-treated fruits of *V7, r, rin*, and *nor*, but not phytoene and phytofluene, similar to CPTA-treated leaves, is seemingly consistent with this view. Ostensibly, CPTA treatment unmasks the PSY2-mediated pathway by terminating it at lycopene. Conceivably PSY2-mediated pathway contributes to the formation of a different set of carotenoids, such as β-carotene, lutein, and violaxanthin, which accumulates in PSY1 as well as in nonripening mutant fruits. Altogether PSY2-mediated pathway operates in parallel but independently of fruit-specific carotenogenesis.

Among the downstream carotenoids β-carotene, lutein, and violaxanthin, the above pathway is more inclined towards the β-carotene branch. Consistent with this, CPTA-treated fruits of *V7, r, rin*, and *nor* mutants show a higher reduction in β-carotene and violaxanthin, while a decline in lutein level is seen only in the *rin* mutant. It is equally plausible that the higher reduction of β-carotene and violaxanthin is due to more effective blockage of the β-carotene branch by CPTA than the α-carotene branch (**La Rocca et al., 2007**), which is also seen in leaves. Such a bias is also apparent by the drastic reduction of β-carotene levels in CPTA-treated IIHR2866 fruits, while CPTA did not affect the lutein level.

Since the carotenoids are also subjected to cleavage by CCDs, leading to volatiles and apocarotenoids (**Hou et al., 2016)**, it is logical to expect a shift in the volatile profiles, specifically those derived from β-carotene. The volatile analysis is consistent with this view as CPTA-treated fruits show a higher abundance of lycopene-derived volatiles and a reduction in β-carotene derived β-ionone. Considering that CCDs expression is not altered in CPTA-treated fruits, the above shift in volatile levels reflects the increase or decrease in respective substrates.

The sustenance of a PSY2 mediated β-carotene synthesis in fruits may be an adaptive means to ensure the accumulation of ABA and other essential apocarotenoids in the developing seeds of fruits. In the above context, the operation of the β-carotene pathway in *V7, r, rin*, and *nor* mutants seem to be essential for sustaining a threshold level of the plant hormone ABA, which is derived from the β-carotene branch (**Xiong and Zhu, 2003**). The reduction in ABA levels in parallel with β-carotene in *V7, r, rin*, and *nor* mutants after CPTA treatments is consistent with this view. This also entails that ABA biosynthesis in tomato fruits is not solely dependent on PSY1-derived carotenoids, and the absence of the PSY1 pathway is compensated by PSY2.

### Accumulation of lycopene triggers chromoplasts-like plastids in *rin* and *nor* mutants

A characteristic marker of fruit ripening in tomato is the conversion of the chloroplasts to chromoplasts, which is triggered in the post-MG phase. The fruits of *V7* and *r* mutants are not blocked in the above transition but lack typical features of tomato chromoplasts such as lycopene threads. Conversely, chromoplast formation is stalled in *rin* and *nor* mutants, as characteristic loss of thylakoids in these mutants is compromised. It is believed that ectopic accumulation of carotenoids in photosynthetic tissues can initiate the differentiation of chromoplast-like plastids. Thus, CPTA-treated leaves show chloroplasts trans-mutating to chromoplasts. Consistent with this, CPTA-treated *rin* and *nor* mutant fruits show loss of thylakoids and the appearance of lycopene threads. Likewise, *V7* and *r*, mutants, where thylakoids dissociate in ripened fruits, after CPTA-treatment, show the appearance of lycopene threads. It can be conceived that the ectopic accumulation of lycopene trigger chromoplast-like plastid in a fashion similar to phytoene-mediated chromoplasts formation in leaves (**Llorente et al., 2020**).

### Accumulation of lycopene in PSY1 knockout mutant is in conformity with operation of independent pathway

It is plausible that in *rin* and *nor* mutants, which have a normal complement of carotenoid biosynthesis genes, in CPTA-treated fruits, the basal expression of *PSY1* may contribute to lycopene accumulation. Likewise, it can also be argued that even in *V7* and *r* mutants, there may be a leaky expression of *PSY1*, which may be responsible for lycopene accumulation. To discount the above possibilities, we used *yellow flesh locus r*^*3756*^ mutant that has a premature stop codon in *PSY1*, terminating the protein at 150 amino acids compared to the wild-type protein of 413 amino acids. The *r*^*3756*^ mutant does not complement the *tangerine* mutant of tomato, retaining the yellow fruit phenotype in *r*^*3756*^/*tangerine* double mutant, indicating that *r*^*3756*^ is a total *PSY1* knockout mutant (**Kachanovsky et al., 2012**). The appearance of lycopene in CPTA-treated *r*^*3756*^ fruits is consistent with the view that either PSY2 or PSY3 contributes to the above response.

The comparison of gene expression profiles of *PSY2* and *PSY3* in tomato fruits favor *PSY2* as the likely contributor. During tomato-fruit ripening, the *PSY1* expression is superinduced post-MG phase, whereas *PSY2* has a low level of consistent constitutive expression right from anthesis till full ripening (**Fantini et al., 2013; Figure S9**). Relative to leaf, the *PSY2* transcript level in ripening fruits (BR+10 days) is only 5-fold lower, whereas *PSY1* shows 57-fold stimulation (**Fantini et al., 2013**). Contrasting to *PSY1* and *PSY2*, the *PSY3* expression is either too low or below the limit of detection during fruit development and ripening. Our qRT-PCR results are consistent with the above analysis; the ripe tomato fruits express the *PSY2* gene, albeit at a much lower level than *PSY1*. The expression of both *PSY1* and *PSY2* is largely unaffected by CPTA-treatment. However, this lower expression of *PSY2* seems to be sufficient to sustain a leaf-like carotenogenesis, which gets uncovered on CPTA-treatment of *psy1* mutants.

### Dark-incubation of isomerization mutants abolishes the CPTA effect

An alternative way to show functional *PSY2* or *PSY3* in tomato fruits is to use a knockout mutant of the carotenogenesis pathway downstream of phytoene synthase. The mutations in carotenogenesis genes are lethal, as blockage of carotenogenesis leads to photooxidation of chloroplasts, barring mutations in CRTISO and ZISO. It is believed that in *CRTISO* and *ZISO* mutants, *cis*-carotenoids intermediates are photoisomerized in the light, thus obviating the need for these enzymes for mutant plants’ survival. However, these enzymes are needed in darkness for carotenoid isomerizations **(Isaacson et al., 2002; Park et al., 2002; Chen et al., 2010**).

Consistent with the above-mentioned light-mediated isomerization, the CPTA-treated *ZISO* mutant fruits incubated in light displayed lycopene accumulation. The reduction of lycopene accumulation in dark-incubated controls (*r* and *r*^*3756*^) and total loss in CPTA-treated *ZISO* mutant fruits is supportive of the notion that lycopene in CPTA-treated fruit is derived from the operation of PDS, CRTISO, ZISO, and ZDS in tomato fruits. The fruits of *yellow oxheart* mutant defective in CRTISO showed very little lycopene accumulation without CPTA treatment, though they accumulate phytoene and phytofluene. Considering that unlike in leaves/seedlings, the light cannot penetrate deep into the fruit tissues, light incubated CRTISO mutant seedlings accumulate the lycopene, but not in the fruits (**Isaacson et al., 2002)**. Nonetheless, the CPTA-treated *yellow oxheart* fruits accumulated lycopene, visible as red tinge in the pericarp periphery of cut sections of fruits, where light likely penetrated.

### Compartmentalization of PSY1 and PSY2 may be different in chromoplasts

Considering that both PSY2 and PSY1 coexist in the fruit, it is difficult to ascribe their relative contribution to phytoene formation in wild-type plants. It is plausible that higher *in vitro* enzymatic formation of phytoene from GGPP in chromoplasts of *r* mutant fruits observed by **Fraser et al. (1999)** arose from residual PSY2 activity. The PSY2 and PSY1 have different Km, pH optima, and cofactor requirements (**Fraser et al., 2000**), thus likely spatially located at different sites in the plastid. Consistent with this, the recent comparisons of tomato PSY2 and PSY1 revealed that compared to PSY1, which has weak enzyme activity, PSY2 is enzymatically more efficient (**Cao et al., 2019**). A more efficient PSY2 can thus sustain carotenogenesis in *PSY1* mutant fruits. Taken together, CPTA-induced lycopene formation in *r* and other carotenogenesis mutants seems to arise from remnant PSY2 activity in ripening fruits. Akin to our results in tomato, in *PSY1*-knockout fruits of Capsicum, a Solanaceae member, PSY2, seems to sustain the basal carotenoid synthesis (**Jang et al., 2020**).

Contrary to the reports that CPTA may enhance the expression of upstream carotenoid biosynthesis genes (**Al-Babili et al., 1999)**, our study did not show any CPTA-specific effect on the expression of both methylerythritol 4-phosphate pathway and carotenoid biosynthesis pathway genes. Ostensibly, CPTA-influence is limited to the inhibition of the enzyme activities of CYC, LCYB1, and LCYB2. The accumulation of lycopene does not lead to a feed-forward stimulation of upstream precursors, as indicated by the absence of phytoene and phytofluene in CPTA-treated fruits of *V7, r, rin*, and *nor* mutants, as well as in CPTA-treated leaves. Conversely, the accumulation of phytoene and phytofluene in fruits having functional PSY1 indicates that PSY1- and PSY2-mediated pathways may be localized in different compartments. Such distinct compartmentalization, not yet reported in tomato, was observed for maize phytoene synthases. In maize plastids, fluorescent protein-tagging revealed that Zm-PSY1 is localized in the stroma, while Zm-PSY2 and Zm-PSY3 were associated with plastoglobule and thylakoids (**Shumskaya et al., 2012**). Moreover, compartmentalization of PSY may differ in a developmental-specific fashion, as potato PSY2 was located in concentrated foci in mesophyll chloroplasts. However, in tuber amyloplasts, it had uniform distribution in the stroma (**Pasare et al., 2013)**.

To sum, our results reveal that while PSY1 is the key enzyme for fruit-specific lycopene and β-carotene accumulation, the ripening fruits in parallel operate another carotenogenesis pathway mediated by PSY2. An important question is why PSY2 mediated pathway does not accumulate phytoene and phytofluene, a feature that is seen in CPTA-treated fruits where PSY1 remains functional, such as in *green flesh*, IIHR2866, *ZISO*, and *yellow oxheart*. We believe since PSY2-mediated carotenogenesis is the mainstay of photosynthetic tissues, the PSY2-pathway is designed to optimize photosynthesis, and prevent any unwanted damage to photosynthesis and ectopic formation of chromoplasts. In Arabidopsis and *Nicotiana benthamiana* leaves the ectopic phytoene accumulation by expressing bacterial crtB (PSY1) in the chloroplast or even extraplastdial compartment interferes with photosynthesis and triggers chromoplast formation (**Llorente et al., 2020; Andersen et al., 2021**).

We presume to prevent photosynthetic damage PSY2-mediated carotenogenesis is carried out by a metabolon different from PSY1-metabolon. Consistent with this in *V7, r*, and *r*^*3756*^ mutants where PSY1 is inactive, and PSY2 is operational, β-carotene and lutein are first accumulated carotenoids, signifying a leaf-like pathway. While this denotes the operation of a pathway independent of PSY1, it also highlights that PSY2 mediated pathway is different and designed to prevent accumulation of potentially photooxidative carotenoid intermediates before β-carotene. Consistent with this notion CPTA-treated leaves also do not show accumulation of phytoene and phytofluene. It remains to be determined how downstream enzymes interact with PSY1 and PSY2 and form two distinct metabolons. At the functional level, these two pathways are ostensibly designed to perform distinct tasks. The PSY1-pathway is geared towards producing attractants for frugivores, the PSY2-pathway is backup for sustaining the essential need of providing abscisic acid during fruit development and ripening.

## Materials and Methods

### Plant material and growth conditions

Seeds of tomato (*Solanum lycopersicum*) cultivar Arka Vikas (AV) and IIHR 2866 were obtained from IIHR (Indian Institute of Horticulture Research), Bangalore. The *rin* (LA1795), *nor* (LA3770), *r* (LA2997), and *gf* (LA3534) mutants were obtained from the Tomato Genetics Resource Center (University of California, Davis, CA), *V7* heirloom from Amishlands Heirlooms, USA, *Yellow Oxheart* heirloom from Victory seeds, Oregon, USA, and *r*^*3756*^ seeds was the generous gift from Dr. Joseph Hirschberg, Hebrew University of Jerusalem, Israel. The *ZISO* mutant was isolated using forward genetics from an EMS remutagenized population (**Gupta et al., 2017**). The tomato fruits used in supplementary figures were obtained from tomatoes grown in an open field sourced from a farmer.

Tomato seeds were surface sterilized with 4% (w/v) NaClO_4_ for 10 min, followed by washing in running tap water and sowing on germination paper. The germinated seedlings were transferred to 50-well nursery trays filled with coco peat (Sri Balaji Agroservices, Madanapalle, Andhra Pradesh, India). The trays were kept ie a greenhouse (12-14 h of light, 28±1°C during the day, ambient in the night), and after three weeks, the seedlings were transferred to the pots.

For fruit studies, different concentration of 2-(4-chlorophenylthio)triethylamine hydrochloride (CPTA) was directly injected in the fruits harvested at different stages of ripening. For leaf assays, tomato seedlings were irrigated with CPTA solution.

### Estimation of Carotenoids

The carotenoid analysis was done as described by **Gupta et al. (2015)**. *E. coli* strains containing pAC-BETA-At (Addgene plasmid no. #53288), pAC-ZETA (Addgene plasmid no. #53316), pAC-EPSILON (Addgene plasmid no. #53276), pAC-LYC (Addgene plasmid no. #53270), pAC-85b (Addgene plasmid no. #53282) was a gift from Francis X Cunningham Jr. Growth conditions and carotenoid extraction from bacteria were as described in **Cunningham and Gantt (2005)**.

### Quantitative Real-Time PCR

Total RNA was extracted from the pericarp tissue of fruits using TRI reagent (Sigma-Aldrich) according to the manufacturer’s protocol. The isolated RNA was incubated with RNAse-free DNAse (Promega) as per the manufacturer’s protocol to eliminate any genomic DNA contamination. The cDNA was prepared from 2 µg RNA using a cDNA synthesis kit (SuperScript III; Invitrogen, USA). Quantitative real-time PCR (qRT-PCR) was carried out in the Aria-MX real-time PCR system (Agilent Technologies). The transcript abundance was measured in 10 µL volume of the SYBR Green PCR Master Mix (Takara, Japan) containing cDNA corresponding to 5 ng of total RNA with gene-specific primers. The ΔCt value was calculated by normalizing each gene’s Ct values to the mean expression of the two internal control genes (β*-actin* and *ubiquitin3*). The list of primers is given in **Table S1**.

### Volatile analysis

The volatile extraction, separation on GC-MS, and identification were carried out as described previously (**Kilambi et al., 2021**). The list of identified volatiles is given in Table S2.

### Plastid Ultrastructure

The pericarp tissue from tomato fruits was excised into approximately 1 mm^3^ pieces and stored in 2.5% glutaraldehyde in 0.1 M cacodylate buffer until further use. Fixed samples were washed with buffer and post-fixed with 1% OsO_4_ in water at 4°C for 2 h. After washing, samples were incubated in 1% Ur Ac for 2 h in darkness and dehydrated with EtOH graded series from 30% to 100 % absolute EtOH. Embedding in epoxy-resin was gradual, using 3:1, 1:1, and 1:3 EtOH/Epon resin ratios (each 12 h) before incubating the samples for 3 days in pure Epon resin, with changes of resin every 12 h. Samples were transferred to molds with fresh resin and let polymerize for 2 days at 60°C. Ultra-thin sections of approximately 70 nm were obtained with a Reichert-Jung ultramicrotome. Sections were collected in 100 mesh copper grids and post-stained with 1% Ur Ac and lead citrate, according to **Reynolds (1963)**.

## Abbreviations

CPTA: 2-(4-Chlorophenylthio) triethylamine hydrochloride
CrtISO: carotenoid isomerase
CYCB: chromoplast-specific lycopene-β-cyclase
IG: immature-green
MG: mature-green
LCYB: chloroplast-specific lycopene β-cyclases
LCYE: lycopene ε-cyclase
PDS: phytoene desaturase
PSY: phytoene synthase
ZISO: ζ-carotene isomerase
ZDS: ζ-carotene desaturase.

## Acknowledgments

This work was supported by the Department of Biotechnology (DBT), India grants, BT/PR11671/PBD/16/828/2008, BT/PR/7002/PBD/16/1009/2012, and BT/COE/34/SP15209/2015 to RS and YS. RS gratefully acknowledges Alexander Von Humboldt’s support for a collaborative visit to Marta Rodriguez□Franco laboratory at the University of Freiburg, Germany. We thank Rosula Hinnenberg for excellent technical assistance during TEM sample preparation.

## AUTHOR CONTRIBUTIONS

PG, YS, and RS designed this project and wrote the manuscript. PG performed most of the experiments. MRF did the electron microscope imaging. RB assisted in the isolation of *ziso* mutant. All authors read and approved the manuscript.

## Data Availability

All data associated with this manuscript are provided in the main paper and supplemental data.

## Supplemental Information

Supplemental tables and figures are available at Online

## Supplemental Data

**Figure S1:** Inhibitory effect of CPTA on the accumulation of lycopene in carotenoid-producing *E. coli* strains

**Figure S2**: Optimization of CPTA concentration for injection.

**Figure S3:** CPTA-induced pigmentation in tomato fruits of different maturity.

**Figure S4:** CPTA-induced lycopene thread-like structures in plastids.

**Figure S4:** Progressive color development in CPTA-treated and control fruits.

**Figure S6:** Carotenoid profiling of CPTA- and water-treated fruits at different days post-injection.

**Figure S7:** Transcript levels of carotenoid pathway genes in *V7, r*, IIHR 2866, *rin, nor*, and *gf* fruits injected with water or CPTA at MG stage.

**Figure S8:** Precocious appearance of carotenoids in plastids of CPTA-injected mutant fruits.

**Figure S9**: Normalized expression (FPKM) of phytoene synthase genes in different organs of tomato.

**Table S1**: List of the genes and the primer sequences used for RT-PCR analysis.

**Table S2**: Volatiles detected in *V7, r*, IIHR 2866, *rin, nor*, and *gf* fruits injected with CPTA at MG stage.

## References

Al-Babili S, Hartung W, Kleinig H, Beyer P. 1999. CPTA modulates levels of carotenogenic proteins and their mRNAs and affects carotenoid and ABA content as well as chromoplast structure in Narcissus pseudonarcissus flowers. Plant Biology 1, 607–612.

Andersen TB, Llorente B, Morelli L, Torres□Montilla S, Bordanaba□Florit G, Espinosa FA, Rodriguez□Goberna MR, Campos N, Olmedilla□Alonso B, Llansola□Portoles MJ, Pascal AA, Rodriguez-Concepcion M. (2021) An engineered extraplastidial pathway for carotenoid biofortification of leaves. Plant Biotechnology 19:1008.

Baldermann S, Kato M, Kurosawa M, Kurobayashi Y, Fujita A, Fleischmann P, Watanabe N. 2010. Functional characterization of a carotenoid cleavage dioxygenase 1 and its relation to the carotenoid accumulation and volatile emission during the floral development of Osmanthus fragrans Lour. Journal of Experimental Botany 61, 2967–2977.

Barry CS. 2014. Ripening mutants. In “Fruit Ripening: Physiology, Signalling, and Genomics” Eds. Nath P, Bouzayen M, Mattoo AK, Pech JC, CABI, pp 246–258.

Barry CS, McQuinn RP, Chung MY, Besuden A, Giovannoni JJ. 2008. Amino acid substitutions in homologs of the STAY-GREEN protein are responsible for the green-flesh and chlorophyll retainer mutations of tomato and pepper. Plant Physiology 147, 179–187.

Camagna M, Grundmann A, Bär C, Koschmieder J, Beyer P, Welsch R. 2019 Enzyme fusion removes competition for geranylgeranyl diphosphate in carotenogenesis. Plant Physiology. 179, 1013–1027

Campisi L, Fambrini M, Michelotti V, Salvini M, Giuntini D, Pugliesi C. 2006. Phytoene accumulation in sunflower decreases the transcript levels of the phytoene synthase gene. Plant Growth Regulation 48, 79–87.

Cao H, Luo H, Yuan H, Eissa MA, Thannhauser TW, Welsch R, Hao YJ, Cheng L, Li L. 2019. A neighboring aromatic-aromatic amino acid combination governs activity divergence between tomato phytoene synthases. Plant Physiology 180, 1988–2003.

Chamovitz D, Pecker I, Hirschberg J. 1991. The molecular basis of resistance to the herbicide norflurazon. Plant Molecular Biology 16, 967–974.

Chen Y, Li F, Wurtzel ET. 2010. Isolation and characterization of the Z-ISO gene encoding a missing component of carotenoid biosynthesis in plants. Plant Physiology 153, 66–79.

Coggins CW, Henning GL, Yokoyama H. 1970. Lycopene accumulation induced by 2-(4-chlorophenylthio)-triethylamine hydrochloride. Science 168, 1589–1590.

Cunningham FX, Gantt E. 2005. A study in scarlet: enzymes of ketocarotenoid biosynthesis in the flowers of Adonis aestivalis. The Plant Journal 41, 478–492.

Cunningham FX, Pogson B, Sun Z, McDonald KA, DellaPenna D, Gantt E. 1996. Functional analysis of the beta and epsilon lycopene cyclase enzymes of Arabidopsis reveals a mechanism for control of cyclic carotenoid formation. The Plant Cell 8, 1613–1626.

Fantini E, Falcone G, Frusciante S, Giliberto L, Giuliano G. 2013. Dissection of tomato lycopene biosynthesis through virus-induced gene silencing. Plant Physiology 163, 986–998.

Fraser PD, Kiano JW, Truesdale MR, Schuch W, Bramley PM. 1999. Phytoene synthase-2 enzyme activity in tomato does not contribute to carotenoid synthesis in ripening fruit. Plant Molecular Biology 40, 687–698.

Fraser PD, Romer S, Shipton CA, Mills PB, Kiano JW, Misawa N, Drake RG, Schuch W, Bramley PM. 2002. Evaluation of transgenic tomato plants expressing an additional phytoene synthase in a fruit-specific manner. Proceedings of the National Academy of Sciences USA 99, 1092–1097.

Fraser PD, Schuch W, Bramley PM. 2000. Phytoene synthase from tomato (Lycopersicon esculentum) chloroplasts–partial purification and biochemical properties. Planta 211, 361–369.

Fraser PD, Truesdale M, Bird CR, Schuch W, Bramley PM. 1994. Carotenoid biosynthesis during tomato fruit development. Evidence for tissue-specific gene expression. Plant Physiology 105, 405–413.

Fray RG, Grierson D. 1993. Identification and genetic analysis of normal and mutant phytoene synthase genes of tomato by sequencing, complementation and co-suppression. Plant Molecular Biology 22, 589–602.

Gupta P, Sreelakshmi Y, Sharma R. 2015. A rapid and sensitive method for determination of carotenoids in plant tissues by high-performance liquid chromatography. Plant Methods 11, 5. https://doi.org/10.1186/s13007-015-0051-0

Gupta P, Reddaiah B, Salava H, Upadhyaya P, Tyagi K, Sarma S, Datta S, Malhotra B, Thomas T, Sunkum A, Devulapalli S, Till BJ, Sreelakshmi Y, Sharma R. 2017. NGS-based identification of induced mutations in a doubly mutagenized tomato (Solanum lycopersicum) population. The Plant Journal 92, 495–508

Hou X, Rivers J, León P, McQuinn RP, Pogson BJ. 2016. Synthesis and function of apocarotenoid signals in plants. Trends in Plant Science 21, 792–803.

Inwood W, Yoshihara C, Zalpuri R, Kim KS, Kustu S. 2008. The ultrastructure of a Chlamydomonas reinhardtii mutant strain lacking phytoene synthase resembles that of a colorless alga. Molecular Plant 1, 925–937.

Isaacson T, Ronen G, Zamir D, Hirschberg J. 2002. Cloning of tangerine from tomato reveals a carotenoid isomerase essential for the production of β-carotene and xanthophylls in plants. The Plant Cell 14, 333–342.

Ito Y, Nishizawa-Yokoi A, Endo M, Mikami M, Shima Y, Nakamura N, Kotake-Nara E, Kawasaki S, Toki S. 2017. Re-evaluation of the rin mutation and the role of RIN in the induction of tomato ripening. Nature Plants 3, 866–874.

Jang SJ, Jeong HB, Jung A, Kang MY, Kim S, Ha SH, Kwon JK, Kang BC. 2020. Phytoene Synthase 2 Can Compensate for the Absence of Psy1 in Capsicum Fruit. Journal of Experimental Botany 71, 3417–3427.

Jen JJ, Thomas RL. 1978. Antagonistic effect of CPTA and far-red light on the carotenogenesis in lutescent tomatoes. Journal of Food Biochemistry 2, 23–27.

Kachanovsky DE, Filler S, Isaacson T, Hirschberg J. 2012. Epistasis in tomato color mutations involves regulation of phytoene synthase 1 expression by cis-carotenoids. Proceedings of the National Academy of Sciences USA 109, 19021–19026.

Kavitha P, Shivashankara KS, Rao VK, Sadashiva AT, Ravishankar KV, Sathish GJ. 2014. Genotypic variability for antioxidant and quality parameters among tomato cultivars, hybrids, cherry tomatoes and wild species. Journal of the Science of Food and Agriculture 94, 993–999.

Kilambi HV, Dindu A, Sharma K, Nizampatnam NR, Gupta N, Thazath NP, Dhanya AJ, Tyagi K, Sharma S, Kumar S, Sharma R, Sreelakshmi Y. 2021. The new kid on the block: A dominant-negative mutation of phototropin1 enhances carotenoid content in tomato fruits. The Plant Journal https://doi.org/10.1111/tpj.15206

Kilambi HV, Kumar R, Sharma R, Sreelakshmi Y. 2013. Chromoplast-specific carotenoid-associated protein appears to be important for enhanced accumulation of carotenoids in hp1 tomato fruits. Plant Physiology 161, 2085–2101.

Knypl J. 1969. Inhibition of chlorophyll synthesis by growth retardants and coumarin, and its reversal by potassium. Nature 224, 1025–1026.

La Rocca N, Rascio N, Oster U, Rüdiger W. 2007. Inhibition of lycopene cyclase results in accumulation of chlorophyll precursors. Planta 225, 1019–1029.

Lewinsohn E, Sitrit Y, Bar E, Azulay Y, Ibdah M, Meir A, Yosef E, Zamir D, Tadmor Y. 2005. Not just colors—carotenoid degradation as a link between pigmentation and aroma in tomato and watermelon fruit. Trends in Food Science and Technology 16, 407–415.

Lois LM, Rodríguez-Concepción M, Gallego F, Campos N, Boronat A. 2000. Carotenoid biosynthesis during tomato fruit development: regulatory role of 1□deoxy□D□xylulose 5□phosphate synthase. The Plant Journal 22, 503–513.

Llorente B, Torres-Montilla S, Morelli L, Florez-Sarasa I, Matus JT, Ezquerro M, D’andrea L, Houhou F, Majer E, Picó B, Cebolla J, Troncoso A, Fernie AR, José-Antonio Daròs JA, Rodriguez-Concepcion M (2020) Synthetic conversion of leaf chloroplasts into carotenoid-rich plastids reveals mechanistic basis of natural chromoplast development. Proceedings of the National Academy of Sciences. 117, 21796–803.

Mickey H. 2013. Characterization of Polymorphism in Chromoplast Specific Phytoene Synthase Gene of Tomato. Ph.D. thesis. University of Hyderabad http://hdl.handle.net/10603/214835

Mohan V, Pandey A, Sreelakshmi Y, Sharma R. 2016. Neofunctionalization of chromoplast specific lycopene beta cyclase gene (CYC-B) in tomato clade. PloS one 11(4), e0153333.https://doi.org/10.1371/journal.pone.0153333

Müller P, Li XP, Niyogi KK. 2001. Non-photochemical quenching. A response to excess light energy. Plant Physiology 125, 1558–1566.

Niyogi KK. 1999. Photoprotection revisited: genetic and molecular approaches. Annual Review of Plant Biology 50, 333–359.

Oelmüller R, Mohr H. 1986. Photooxidative destruction of chloroplasts and its consequences for expression of nuclear genes. Planta 167, 106–113.

Park H, Kreunen SS, Cuttriss AJ, DellaPenna D, Pogson BJ. 2002. Identification of the carotenoid isomerase provides insight into carotenoid biosynthesis, prolamellar body formation, and photomorphogenesis. The Plant Cell 14, 321–332.

Pasare S, Wright K, Campbell R, Morris W, Ducreux L, Chapman S, Bramley P, Fraser P, Roberts A, Taylor M. 2013, The sub-cellular localisation of the potato (Solanum tuberosum L.) carotenoid biosynthetic enzymes, CrtRb2 and PSY2. Protoplasma. 250, 1381–1392.

Poling SM, Hsu WJ, Yokoyama H. 1975. Structure-activity relationships of chemical inducers of carotenoid biosynthesis. Phytochemistry 14, 1933–1938.

Reynolds ES. 1963. The use of lead citrate at high pH as an electron-opaque stain in electron microscopy. Journal of Cell Biology 17, 208–212.

Robert B, Horton P, Pascal AA, Ruban AV. 2004. Insights into the molecular dynamics of plant light-harvesting proteins in vivo. Trends in Plant Science 9, 385–390.

Römer S, Fraser PD, Kiano JW, Shipton CA, Misawa N, Schuch W, Bramley PM. 2000. Elevation of the provitamin A content of transgenic tomato plants. Nature Biotechnology 18, 666–669.

Schwartz SJ, von Elbe JH, Giusti MM. 2008. Colorants. In: Damodaran S, Parkin KL, Fennema OR, editors. Fennema’s Food Chemistry. 4th ed. Boca Raton, FL: CRC Press, 571–638.

Shapira OR, Khadka S, Israeli Y, Shani UR, Schwartz A. 2009. Functional anatomy controls ion distribution in banana leaves: significance of Na+ seclusion at the leaf margins. Plant, Cell and Environment 32, 476–485.

Shumskaya M, Bradbury LM, Monaco RR, Wurtzel ET. 2012. Plastid localization of the key carotenoid enzyme phytoene synthase is altered by isozyme, allelic variation, and activity. The Plant Cell 24, 3725–3741.

Simkin AJ, Breitenbach J, Kuntz M, Sandmann G. 2000. In vitro and in situ inhibition of carotenoid biosynthesis in Capsicum annuum by bleaching herbicides. Journal of Agricultural and Food Chemistry 48, 4676–4680.

Simkin AJ, Zhu C, Kuntz M, Sandmann G. 2003. Light-dark regulation of carotenoid biosynthesis in pepper (Capsicum annuum) leaves. Journal of Plant Physiology 160, 439–443.

Spurgeon SL. 1983. Biosynthesis of carotenoids. Biosynthesis of Isoprenoid Compounds 2, 1–122. John Wiley & Sons

Sun T, Yuan H, Cao H, Yazdani M, Tadmor Y, Li Li. 2018. Carotenoid metabolism in plants: the role of plastids. Molecular Plant 11, 58–74.

Thompson AJ, Tor M, Barry CS, Vrebalov J, Orfila C, Jarvis MC, Giovannoni JJ, Grierson D, Seymour GB. 1999. Molecular and genetic characterization of a novel pleiotropic tomato-ripening mutant. Plant Physiology 120, 383–390.

Walter MH, Strack D. 2011. Carotenoids and their cleavage products: biosynthesis and functions. Natural Product Reports 28, 663–692.

Xiong L, Zhu JK. 2003. Regulation of abscisic acid biosynthesis. Plant Physiology 133, 29–36.

Young AJ, Britton G, Musker D. 1989. A rapid method for the analysis of the mode of action of bleaching herbicides. Pesticide Biochemistry and Physiology 35, 244–250.

Zhang Y, Fernie AR. 2020. Metabolons, enzyme-enzyme assemblies that mediate substrate channeling, and their roles in plant metabolism. Plant Communications 2, 100081 https://doi.org/10.1016/j.xplc.2020.100081

