## Supplementary material for "Phytoene synthase 2 in tomato fruits remains functional and contributes to abscisic acid formation": Figure S1

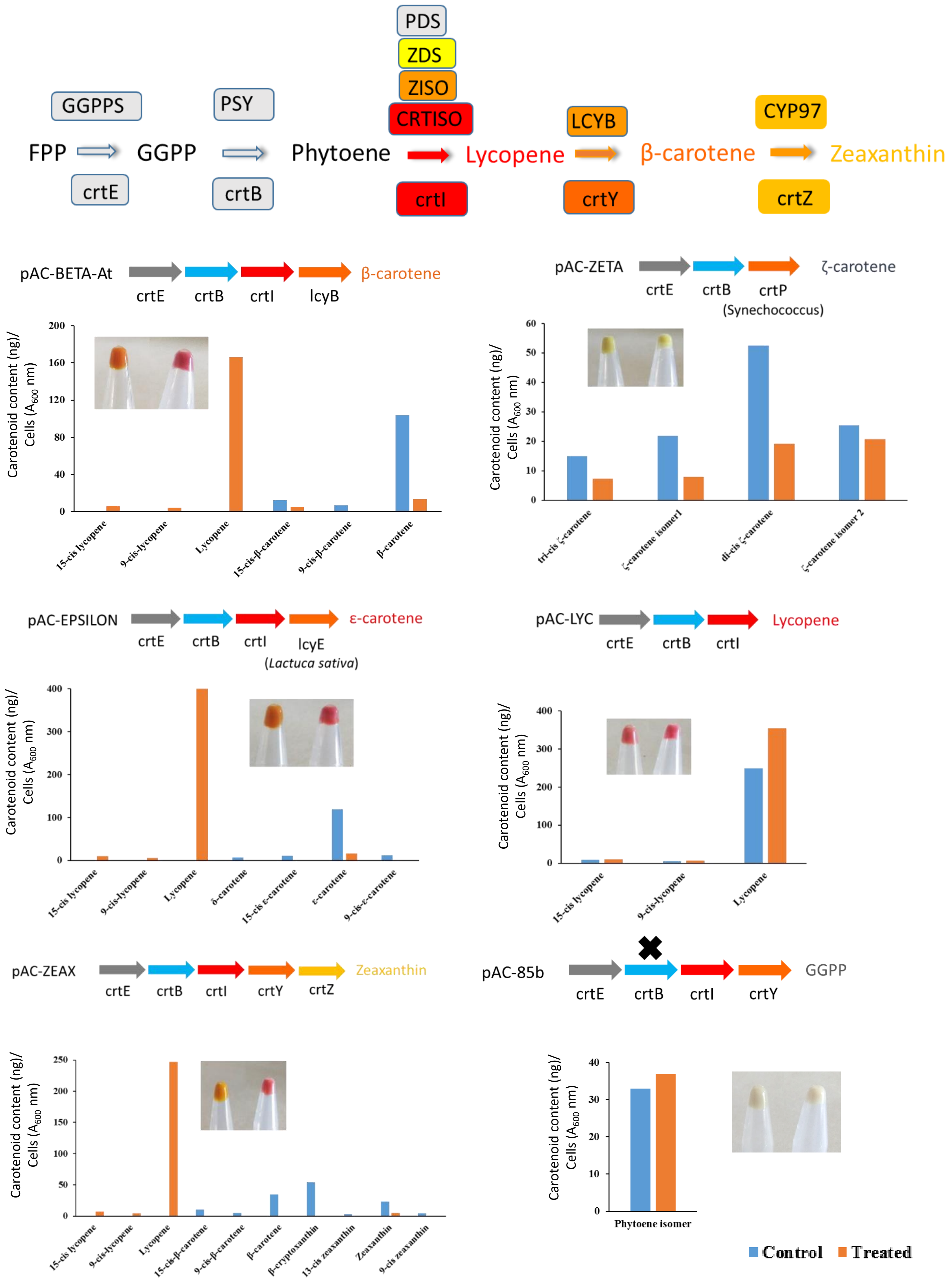


**Figure S1:** Inhibitory effect of CPTA on the accumulation of lycopene in carotenoid-producing *E. coli* strains. The addition of CPTA (1 mM) to pAC-EPSILON, pAC-ZEAX, and pAC-Beta-At produces lycopene as the major carotenoid by inhibiting *LCYE* and *LCYB*, respectively. The addition of CPTA to the pAC-ZETA, pAC-85b, and pAC-LYC does not affect the carotenoid content. The inset picture in the respective graph shows carotenoid accumulation in the bacterial pellet (**left**-Control, **right**-Treated).

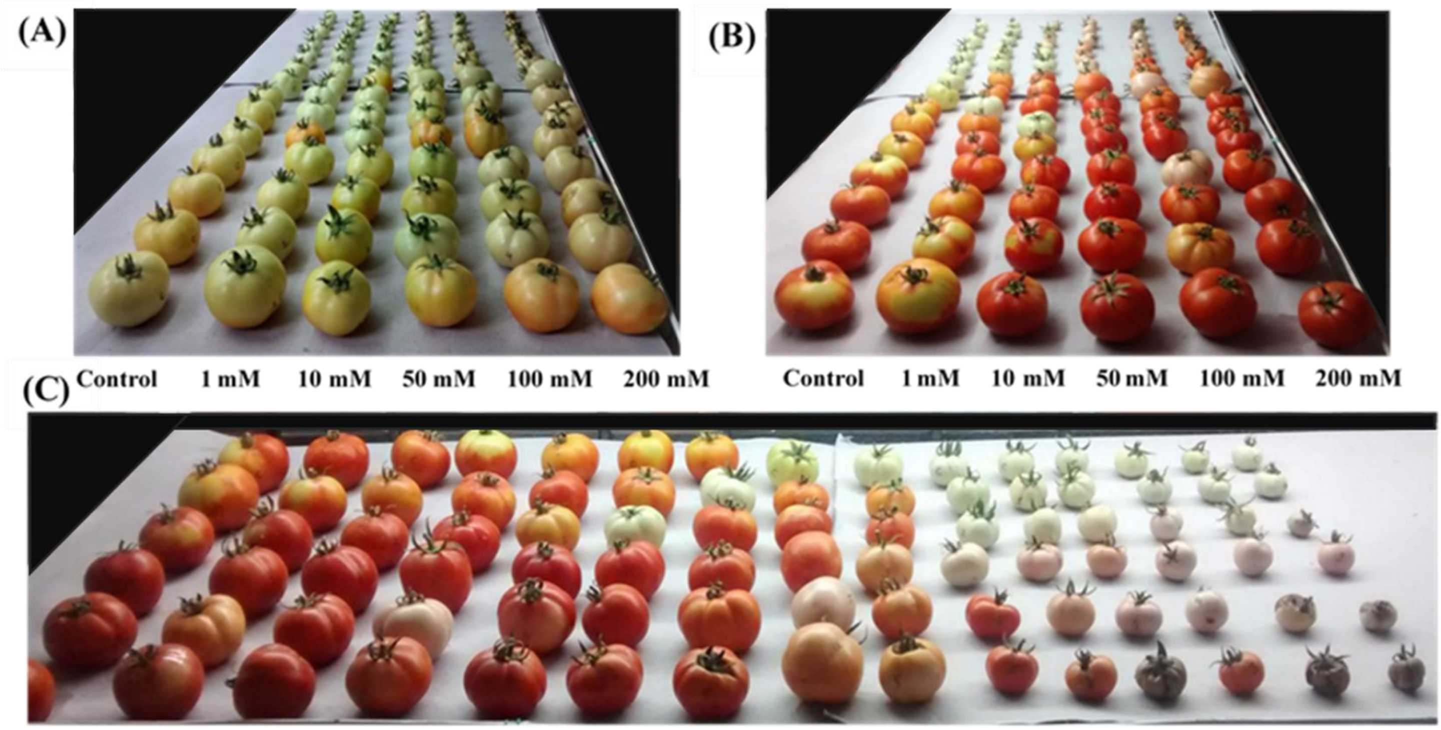


**Figure S2**: Optimization of CPTA concentration for injection. Tomato fruits of different maturity stages from anthesis were used to study the effect of CPTA. Five different concentrations of CPTA (1, 10, 50, 100, and 200 mM) were injected into the detached fruits. The control fruits were injected with water. The fruits were incubated at 25°C under 16h day and 8h dark cycle. The fruits were photographed after three days (**A**) (Side view) and 15 days of injection (**B**) (Side view) and C (Front view). Fruits collected from the farmer's field were used for data in Figure S1 to S5.

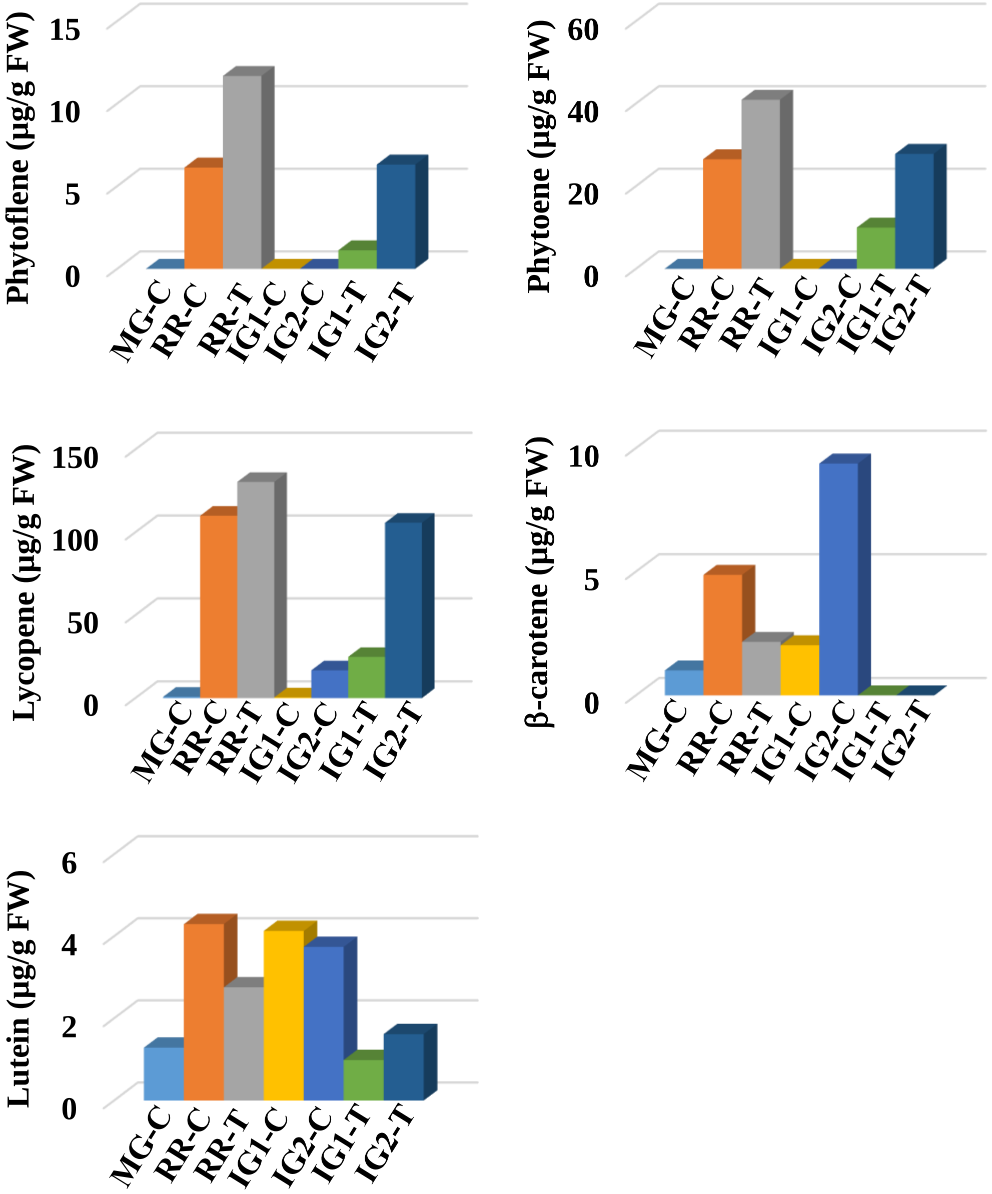


**Figure S3:** CPTA-induced pigmentation in tomato fruits of different maturity. The fruits at different post-anthesis ages (Immature green (**IG**)- IG1- (green color fruit), IG2- (orange color fruit), mature green (**MG**) were injected with CPTA/water. After 13 days of CPTA/water-injection, the carotenoid profile was examined from a single fruit. The fruits post-CPTA injection were incubated under 16h light/8h dark cycles (**C**-Control, **T**-Treated). For fruits with CPTA at MG stage carotenoids levels were determined without injecting CPTA at 0 time (MG-C) and after 13-days with water-injection (RR-C) or with CPTA-injection (RR-T).

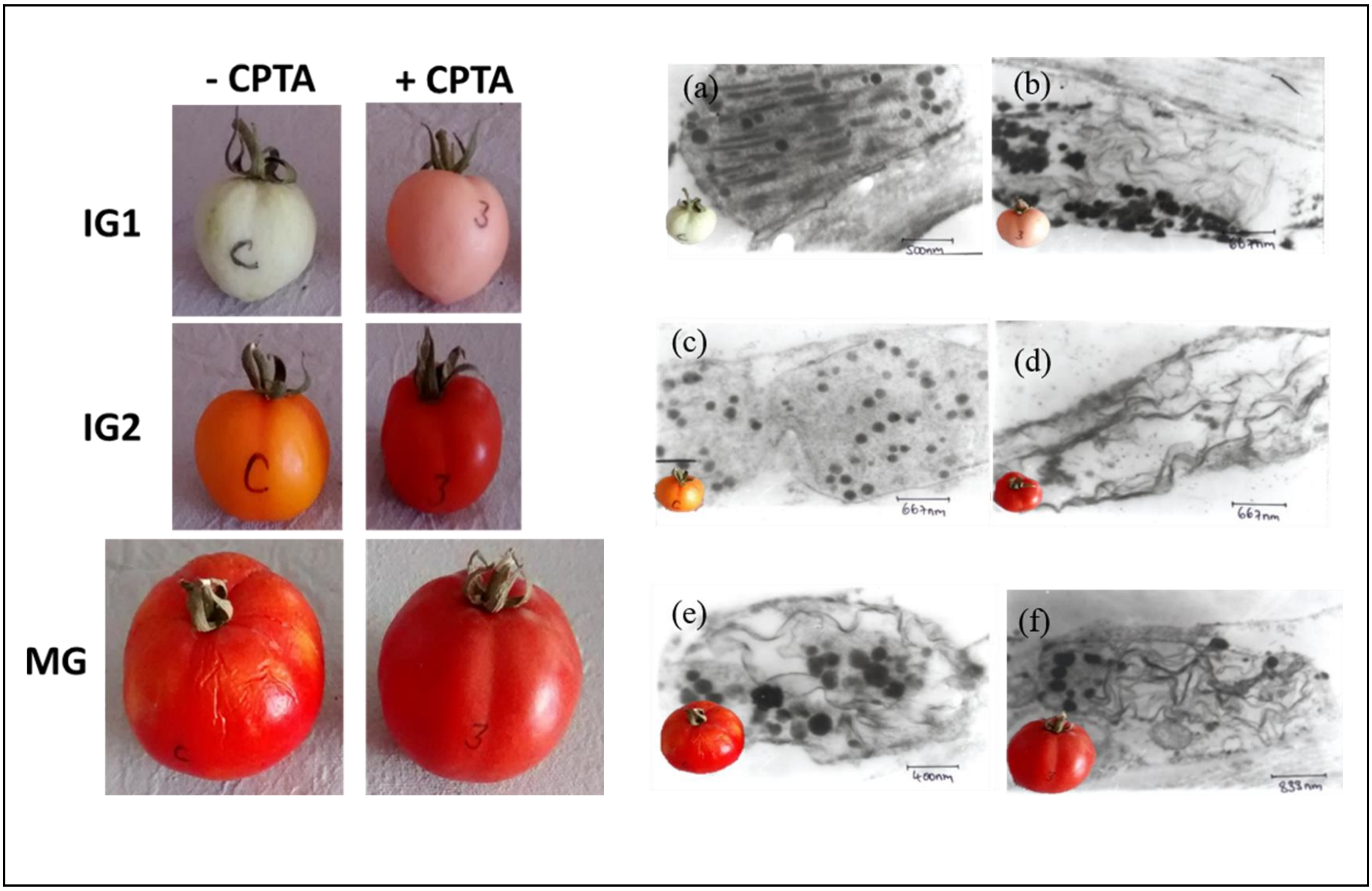


**Figure S4:** CPTA-induced lycopene thread-like structure in plastids. After 13 days of CPTA-treatment, the fruits were photographed, and the ultrastructure of plastids was visualized by TEM. (**a**) IG1 (Control), (**b**) IG1 (Treated), (**c**) IG2 (Control) and (**d**) IG2 (Treated), (**e**) Mature green fruit (Control), (**f**) Mature green fruit (Treated). Note the appearance of precocious lycopene threads in the plastids of IG1 and IG2 fruits. Note the control fruits are shown after 13 days of incubation, during which IG2 fruits accumulate β-carotene precociously.

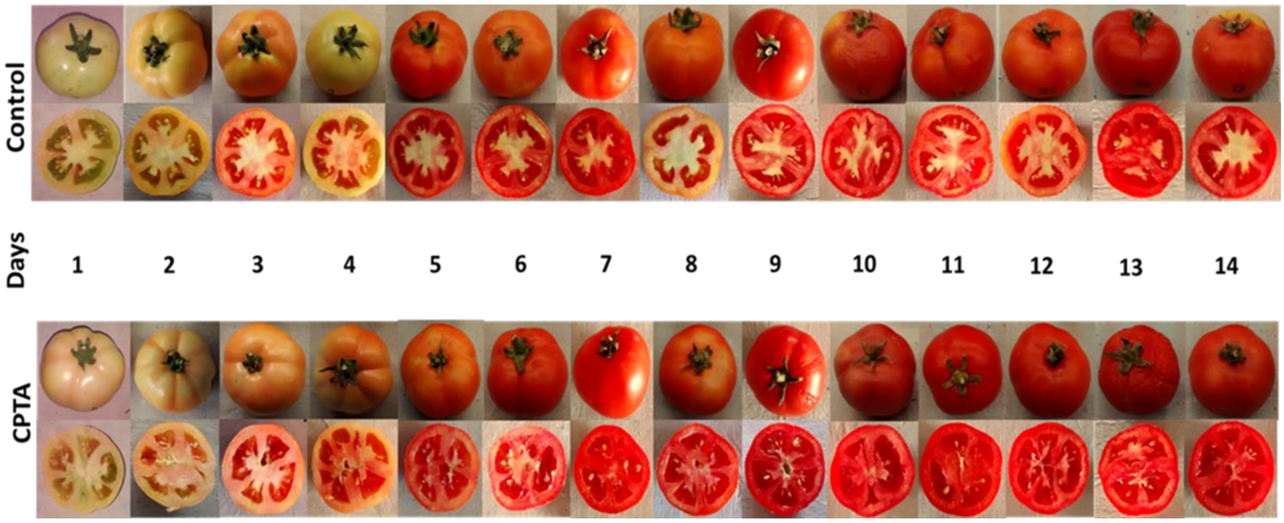


**Figure S5:** Progressive color development in CPTA-treated and control fruits. Note appearance of lycopene in the placenta of CPTA-treated fruits at $4^{th}$-day post-injection and persistence after that. Mature green fruits were injected with 500 µl of 100 mM CPTA, and control fruits were injected with an equal amount of water.

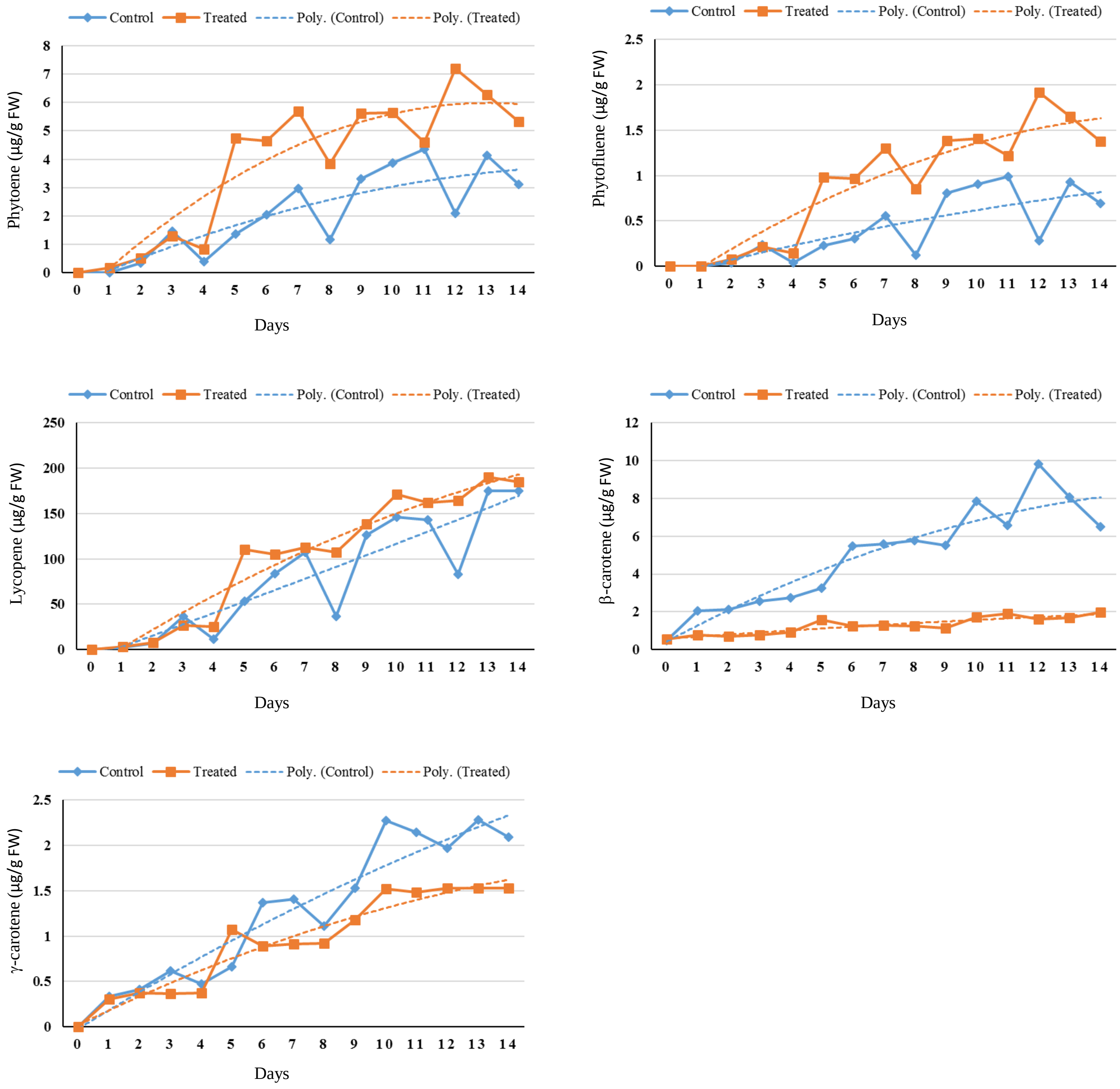


**Figure S6:** Carotenoid profiling of CPTA- and water-treated fruits at different days post-injection. The fruits depicted in Figure S5 were used for the carotenoids profiling (data from single fruit). The dotted line shows the best fit of data using polynomial equation (Order 2) to overcome the variation due to the use of a single fruit. The curve fitting supports increased phytoene, phytofluene, lycopene levels, and the decline of β-carotene level in the treated fruits.

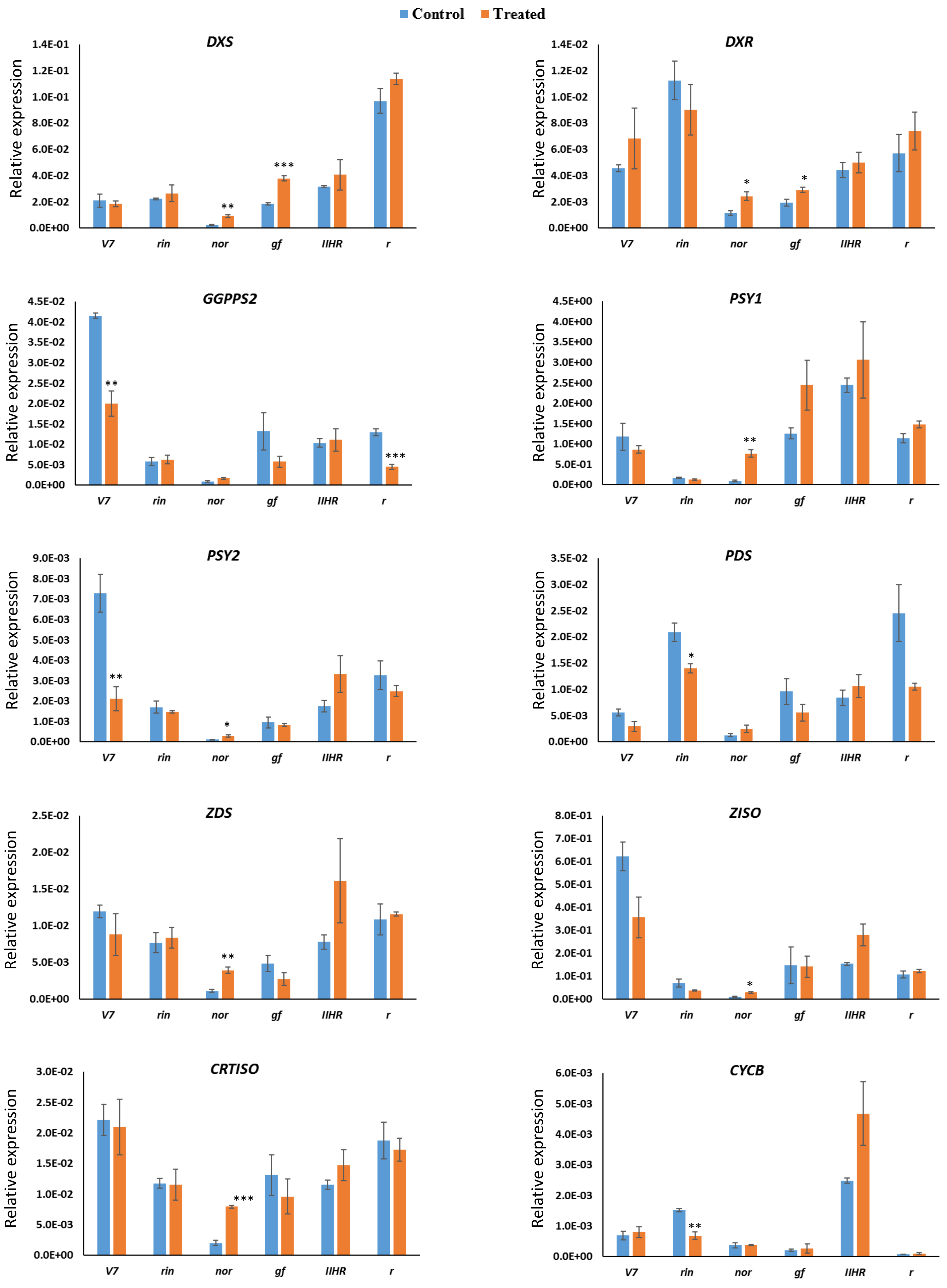
Control
Treated
DXS
DXR
GGPPS2
PSY1
PSY2
PDS
ZDS
ZISO
CRTISO
CYCB
Relative expression
V7
rin
nor
gf
IIHR
r
1.4E-01
1.2E-01
1.0E-01
8.0E-02
6.0E-02
4.0E-02
2.0E-02
0.0E+00
1.4E-02
1.2E-02
1.0E-02
8.0E-03
6.0E-03
4.0E-03
2.0E-03
4.5E-02
4.0E-02
3.5E-02
3.0E-02
2.5E-02
2.0E-02
1.5E-02
1.0E-02
5.0E-03
4.5E+00
4.0E+00
3.5E+00
3.0E+00
2.5E+00
2.0E+00
1.5E+00
1.0E+00
5.0E-01
0.0E+00
9.0E-03
8.0E-03
7.0E-03
6.0E-03
5.0E-03
4.0E-03
3.0E-03
1.0E-03
3.5E-02
3.0E-02
2.5E-02
1.5E-02
2.5E-02
2.0E-02
1.5E-02
1.0E-02
5.0E-03
8.0E-01
7.0E-01
6.0E-01
5.0E-01
4.0E-01
3.0E-01
2.0E-01
1.0E-01
3.0E-02
6.0E-03
5.0E-03
4.0E-03
3.0E-03
2.0E-03
1.0E-03

Cont.....

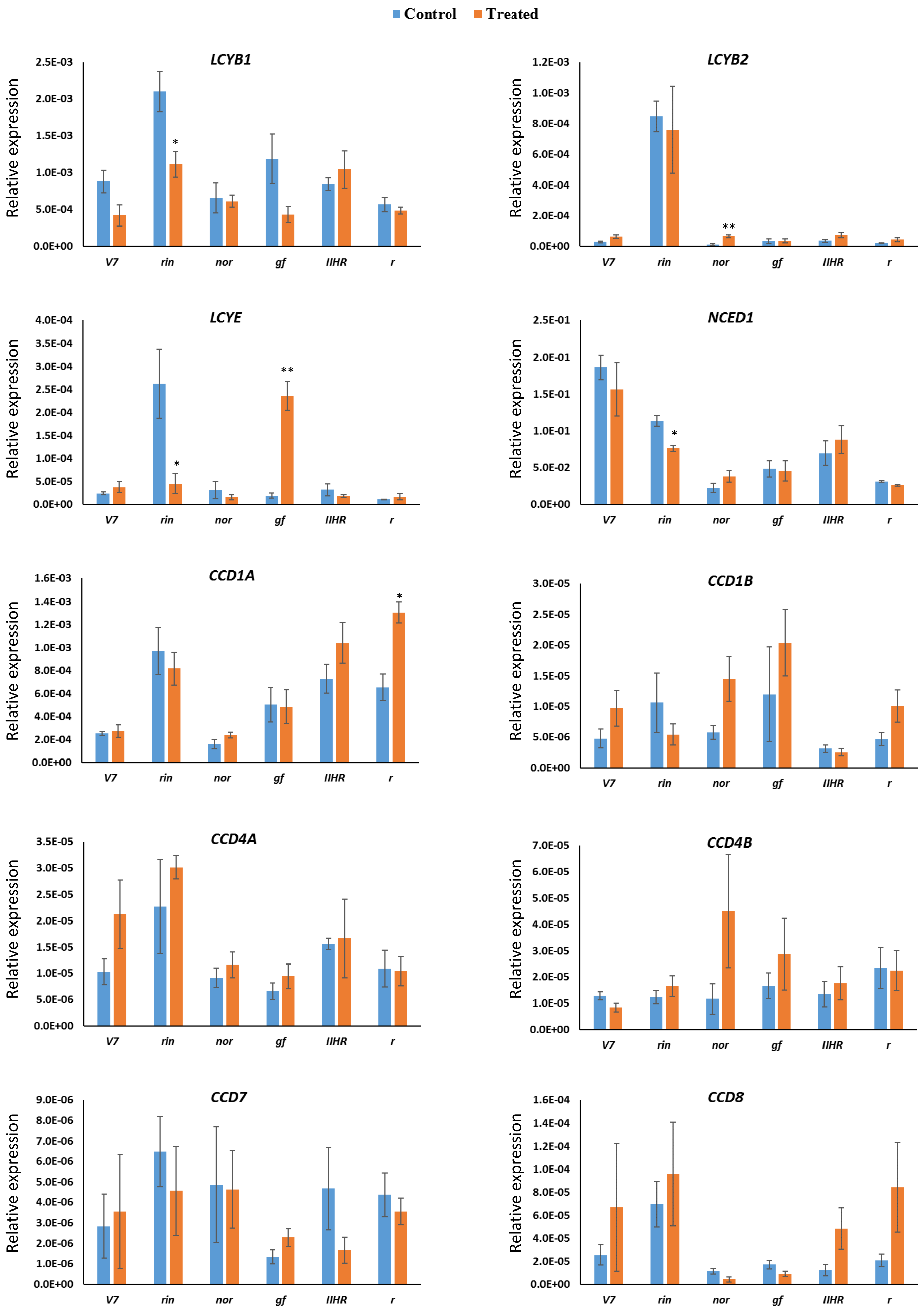


**Figure S7:** Transcript levels of carotenoid pathway genes in *V7*, *r*, IIHR 2866, *rin*, *nor*, and *gf* fruits injected with CPTA at the mature green stage after 12 days of treatment (n>=3, * $p \leq 0.05$, ** $p \leq 0.01$, *** $p \leq 0.001$, p-values are calculated for the treated samples with respect to respective control).

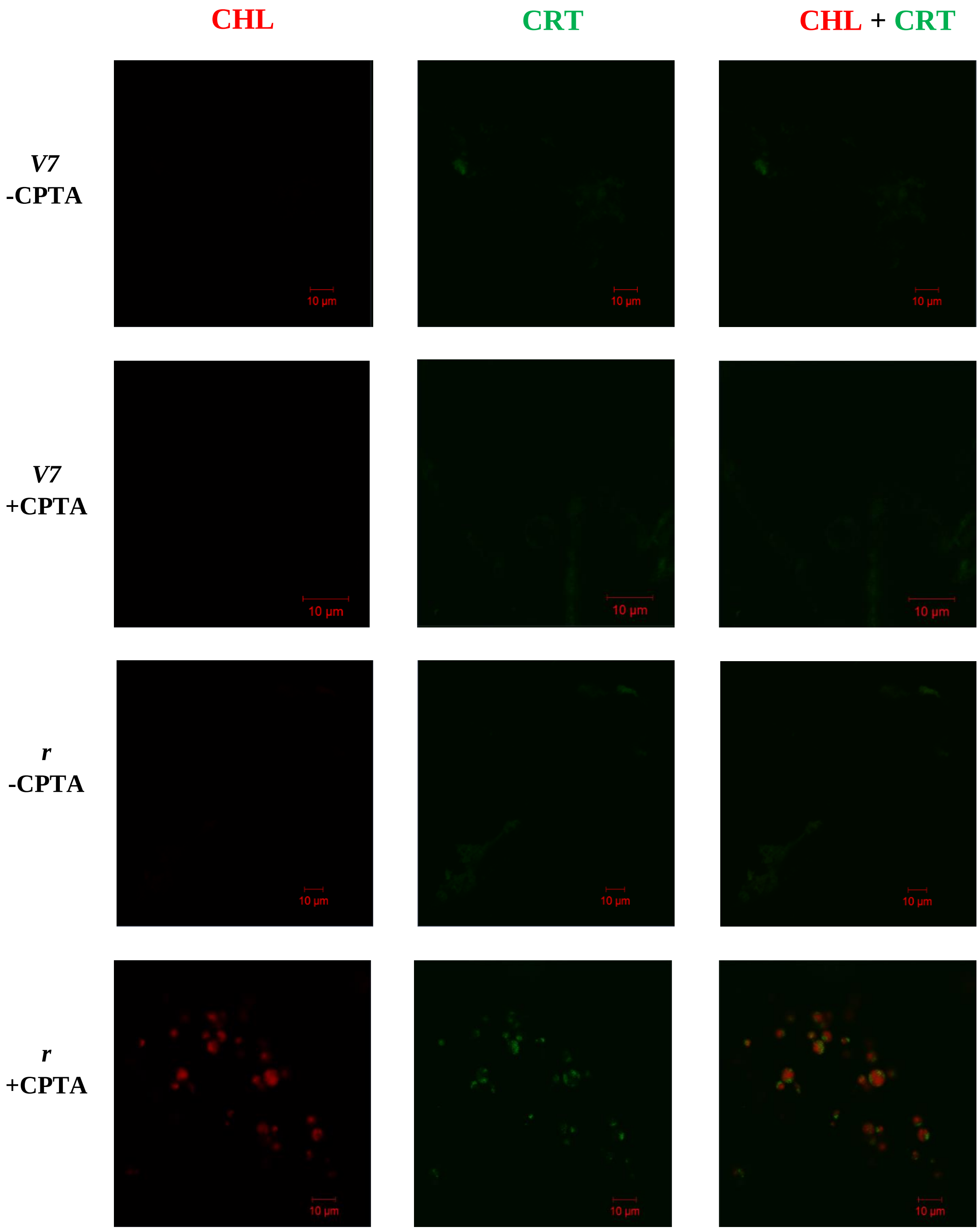


**Cont…..**

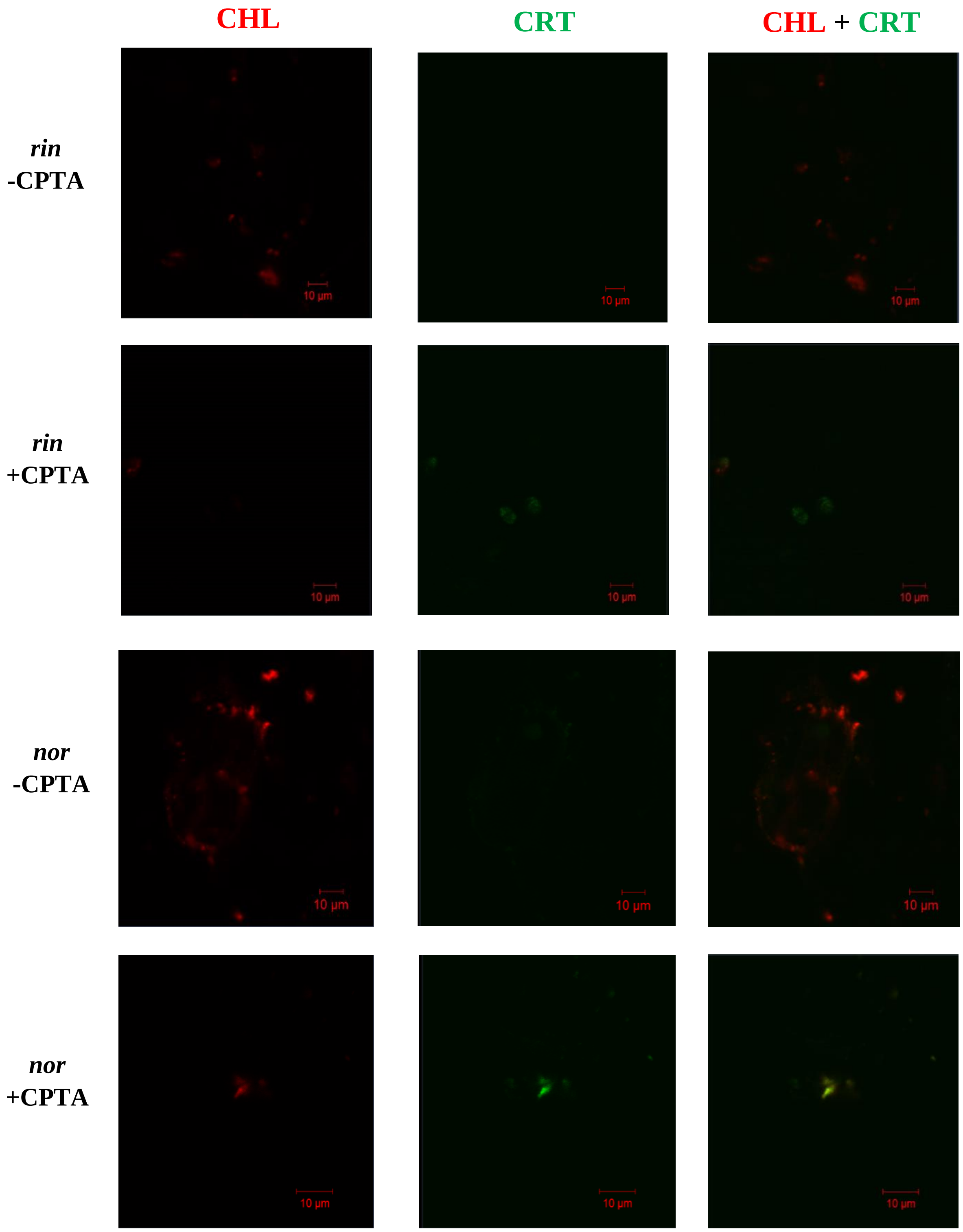


**Figure S8:** Precocious appearance of carotenoids in plastids of CPTA-injected mutant fruits. The freehand sections of the fruit pericarp were examined under a Zeiss confocal microscope after 12-days of CPTA treatment. The sections were observed under the 60X water-immersion objective. The pictures show original and overlaid images of autofluorescence emitted at 650–700 nm (chlorophylls, **CHL**) or 500–550 nm (carotenoids, **CRT**) after excitation with the 488 nm argon laser.

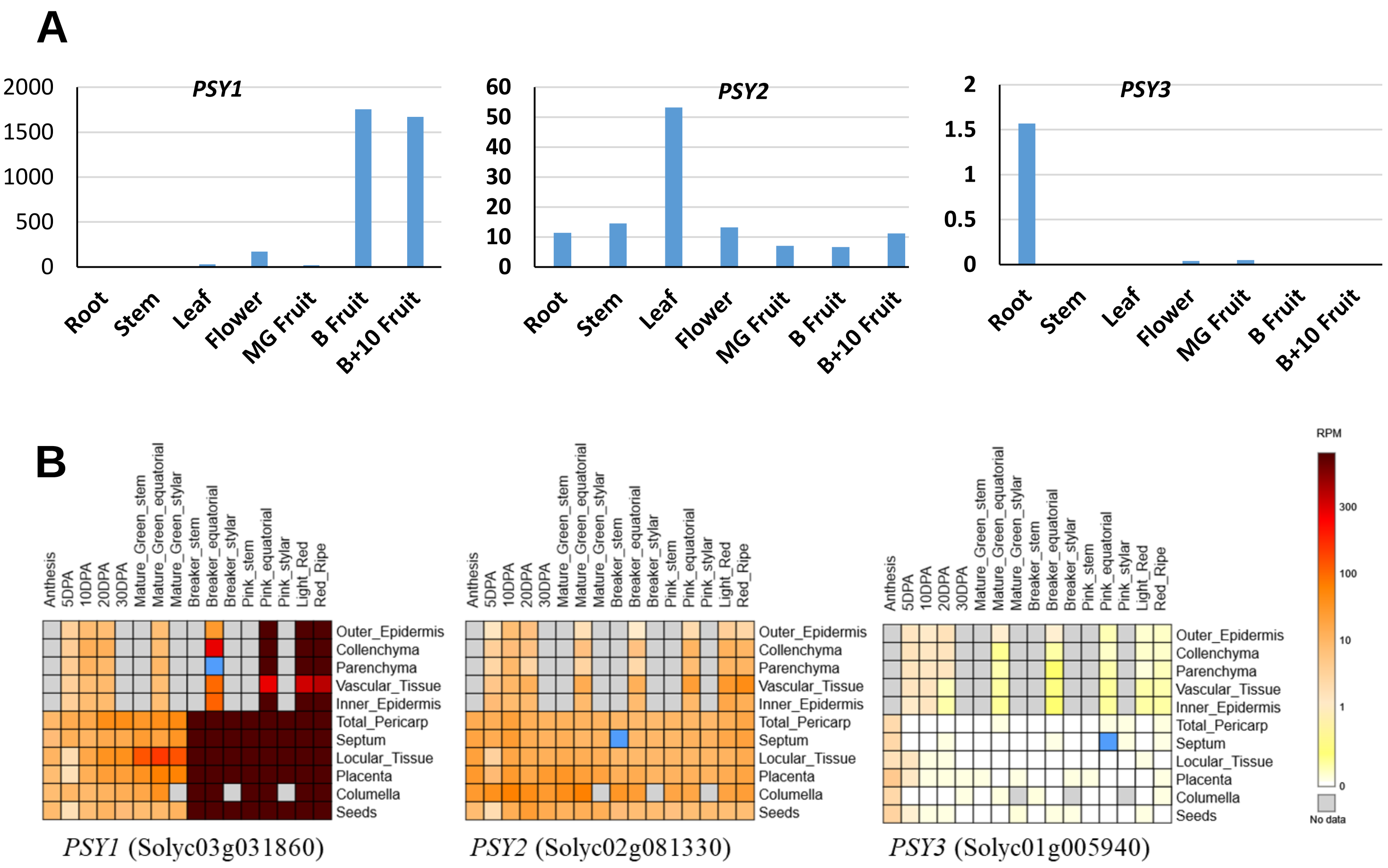


**Figure S9**: The expression of phytoene synthase genes in tomato.

**A.** Normalized expression (FPKM) of phytoene synthase genes in different organs of tomato. **MG**: Mature Green fruit; **B**: Breaker fruit; B+10: ripe fruit 10 days after breaker stage. (Data Source: Fantini et al., (2013) Plant Physiology **163**, 986-998).

**B.** Expression of phytoene synthase genes in tissues of tomato fruits from anthesis to full ripening (Data source https://tea.solgenomics.net/expression_viewer/output).
