## Supplementary material for "Phytoene synthase 2 in tomato fruits remains functional and contributes to abscisic acid formation": Table S1

**Table S1. List of genes and the primers used for qRT-PCR analysis.**

| Gene | Forward primer (5′ to 3′) | Reverse primer (5′ to 3′) | Product size (bp) |
| --- | --- | --- | --- |
| DXS  (Solyc01g067890) | AAATGGGATCGGTGTAGAGC | TGCTGAGCCATATCCCAATA | 115 |
| DXR  (Solyc03g114340) | AGCAGGGTGTGATTGAGGTT | GTCTTTTCCTGCTTCGATGG | 113 |
| GGPPS2  (Solyc04g079960) | ATCAATGGAGCAGCTTTGTG | GCGGTTGATAAAACGACGTA | 128 |
| PSY1  (Solyc03g031860) | TGAATTAGCACAGGCAGGTC | TCAATTCTGTCACGCCTTTC | 140 |
| PSY2  (Solyc02g081330) | AATTCCGAGGTCTCATACGG | CCTTTCCACATCGAATTCCT | 110 |
| PDS  (Solyc03g123760) | TATCATCAACGTTCCGTGCT | TATCGGTTTGTGACCAGCAT | 122 |
| ZDS  (Solyc01g097810) | TCCAAAAGGGCTATTTCCAC | TTGATCCAAGAGCTCCACAG | 115 |
| ZISO  (Solyc12g098710) | AGAGCGTGCTTTTCGTGTATTG | ATTGCCATAACTGCACTCCATC | 107 |
| CRTISO  (Solyc10g081650) | GAGATCGCCAAATCCTTAGC | CAGAAAGCTTCACTCCCACA | 118 |
| CYCB  (Solyc06g074240) | TCTTCTCAAGCCTTTTCCATC | TGGTGGGACTTAGAAAAGAAGG | 92 |
| LCYB1  (Solyc04g040190) | CGATGCAACTGGCTTCTCTA | AATGAGAATCTCGCCAATCC | 149 |
| LCYB2  (Solyc10g079480) | ATTTGTGGCCCATAGAAAGG | TGACAAGAAACCATGCCAAT | 146 |
| LCYE  (Solyc12g008980) | TTAGTCGCCATTTTCTGCAC | TCACCCTCGCACTCTACAAG | 130 |
| NCED1  (Solyc07g056570) | TGACACCACCAGACTCCATT | ACTTGTTCATCCGGGTTTTC | 130 |
| CCD1A  (Solyc01g087250) | TTGACGCATTCCTTCACTGC | GTAAGGTGGGGTGTGAGCAT | 87 |
| CCD1B  (Solyc01g087260) | AGCTAGGAAAATCAAAGGAATAGATGG | TTCAGAAAGAGTGACGTGTTGATC | 107 |
| CCD4A  (Solyc08g075480) | TTCATACTCGCCGGTGGTTC | CACCCTTGGCATAACGTGGA | 85 |
| CCD4B  (Solyc08g075490) | AGACGATGGCTACGTAATGTTG | GGCAATTTAACATTAGCCACAA | 114 |
| CCD7  (Solyc01g090660) | TCTTGCCACCGGCTAAACTG | TTCATGAGTTGGGGACGTGG | 110 |
| CCD8  (Solyc08g066650) | TGTGGTGCTAAGAGGCCTTG | GAATGGTTCAGAAGGCACAGC | 111 |
| β-actin  (FJ532351.1) | TGTCCCTATTTACGAGGGTTATGC | CAGTTAAATCACGACCAGCAAGAT | 108 |
| UBIQUITIN3  (X58253.1) | GCCGACTACAACATCCAGAAGG | TGCAACACAGCGAGCTTAACC | 110 |
